# Direct visualization of four diffusive LexA states controlling SOS response strength during antibiotic treatment

**DOI:** 10.1101/2020.07.14.201889

**Authors:** Leonard Schärfen, Miloš Tišma, Andreas Hartmann, Michael Schlierf

## Abstract

In bacteria, the key mechanism governing mutation, adaptation and survival upon DNA damage is the SOS response. Through autoproteolytic digestion triggered by single-stranded DNA caused by most antibiotics, the transcriptional repressor LexA controls over 50 SOS genes including DNA repair pathways and drivers of mutagenesis. Efforts to inhibit this response and thereby combat antibiotic resistance rely on a broad understanding of its behavior *in vivo*, which is still limited. Here, we develop a single-molecule localization microscopy assay to directly visualize LexA mobility in *Escherichia coli* and monitor the SOS response on the level of transcription factor activity. We identify four diffusive populations and monitor their temporal evolution upon ciprofloxacin-induced continuous DNA damage. With LexA mutants, we assign target bound, non-specifically DNA bound, freely diffusing and cleaved repressors. We develop a strategy to count LexA in fixed cells at different time points after antibiotic stress and combine the time-evolution of LexA sub-populations and the repressor’s overall abundance. Through fitting a detailed kinetic model we obtain *in vivo* synthesis, cleavage and binding rates and determined that the regulatory feedback system reaches a new equilibrium in ∼100 min. LexA concentrations showed non-constant heterogeneity during SOS response and designate LexA expression, and thereby regulation of downstream SOS proteins, as drivers of evolutionary adaptation. Even under low antibiotic stress, we observed a strong SOS response on the LexA level, suggestion that small amounts of antibiotics can trigger adaptation in *E. coli.*

## Introduction

Mechanisms of bacterial resistance to antibiotics include removing drugs *via* various efflux pumps, acquiring resistance genes encoding for drug-inactivating enzymes, or generating mutations in drug target proteins (1). In *Escherichia coli* and many other bacterial species, acquisition of resistance genes or mutations of drug target genes are results of the SOS response, a gene regulatory network that specifically responds to DNA damage, a downstream effect of nearly every antibiotic (2). Despite decades of research, how this regulatory scheme responds to different instances of DNA damage still lacks a global understanding and is hard to predict, in part due to difficulty of monitoring first steps of SOS induction in living cells.

Orchestrated by the transcriptional repressor LexA, over 50 genes involved in DNA repair, mutagenesis, cell cycle control as well as several unknown functions are co-regulated (3). The SOS response is initiated through the exposure of single-stranded DNA (ssDNA), which is a product of nuclease complexes such as RecBCD processing DNA damage (4). Free monomers of RecA are recruited to ssDNA and rapidly polymerize along it (5). The resulting filament can induce autoproteolytic cleavage of the dimeric LexA repressor (6, 7), leading to net dissociation from operator sites and expression of SOS genes such as *sulA, uvrA*, and *umuCD* (3) (Fig. 1a). Since only freely diffusing LexA dimers are susceptible to RecA-mediated autoproteolytic digestion (8), LexA DNA binding kinetics are determinant of the gene expression profile following DNA damage (9). Cleaved LexA fragments are thought to be unstable in the cytoplasm and quickly degraded by multiple proteases recognizing peptide motifs exposed after cleavage (10).

**Figure 1:**
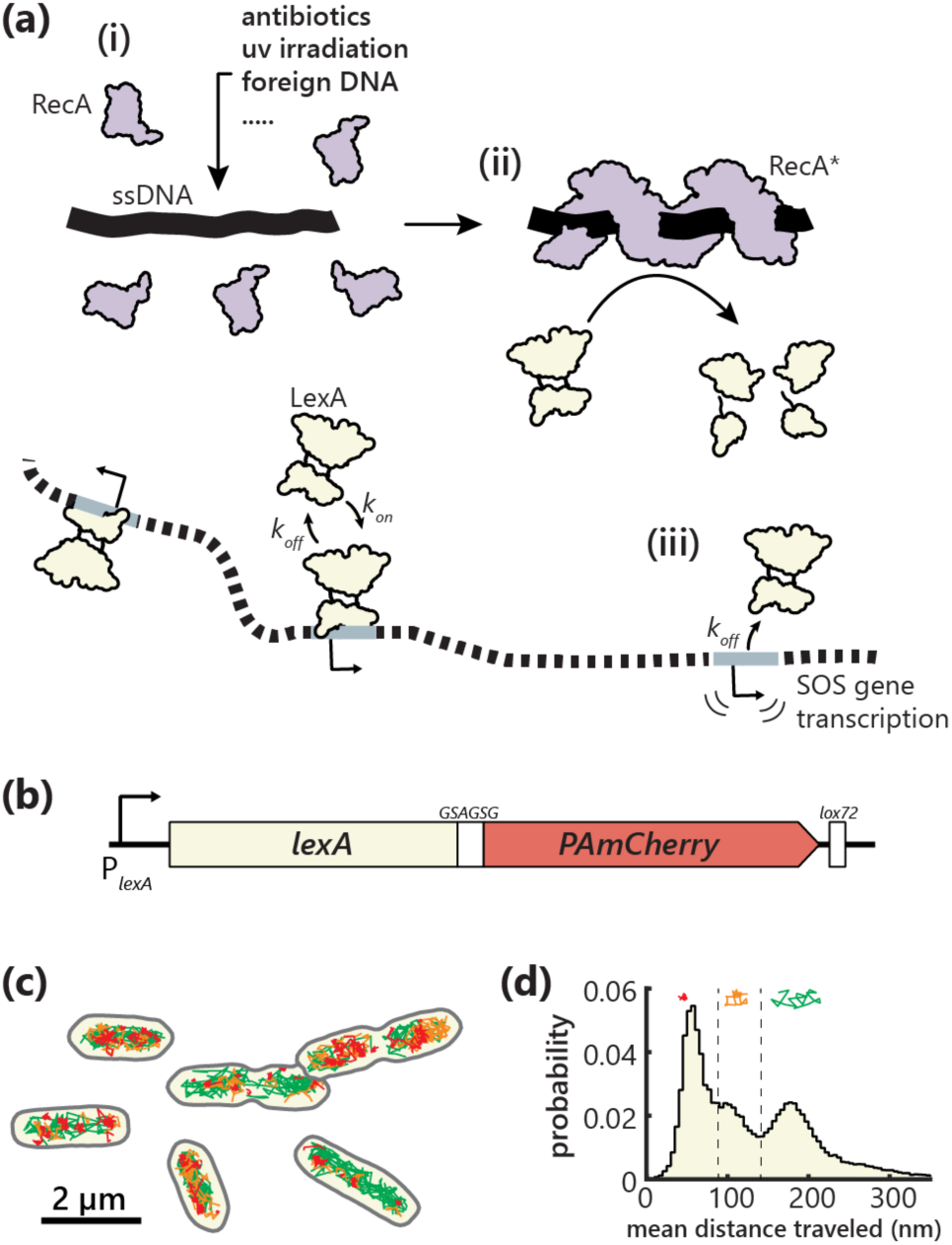
Observing LexA dynamics during SOS response. **(a)** Exposure of single-stranded DNA can be caused by various factors. In unstressed cells, LexA is bound to its target sites and represses SOS genes (i). RecA forms filaments around single-stranded DNA and co-catalyzes LexA autoproteolysis (ii), which leads to less functional LexA dimers in the cytoplasm and a net dissociation from target sites (iii). **(b)** Genomic *lexA* locus after tagging with PAmCherry. **(c)** LexA trajectories in unstressed cells expressing the recombinant protein. Immobile (red), slowly (orange) and fast (green) diffusing molecules, classified by the distance they travel within 20 ms. **(d)** Probability distribution of distance traveled per frame of 73,580 molecules in 1,584 cells.

LexA represses both the *lexA* and *recA* genes in a double negative feedback. Together with the availability of the inducer ssDNA being highly incidental, the expression of SOS genes is nuanced and likely subject to stochastic fluctuations with a high cell-to-cell variability (11, 12). This complexity makes it difficult to predict bacterial behavior under antibiotic challenges. A lack of timescales, molecular quantities and kinetics of individual constituents *in vivo* still blur our understanding of sub-cellular dynamics under stress. Many conventional SOS response assays rely on transcriptional reporters under the control of promoters that become active only after LexA cleavage (9, 11, 13), or on bulk measurements that cannot capture population heterogeneity. More recently, live-cell single-molecule microscopy has developed into a powerful tool in bacteriology because it addresses spatial and temporal scales appropriate for the small size of bacteria (14, 15). Subpopulations of diffusing species can be resolved, and kinetic rates extracted from time course experiments. Fixed-cell single-molecule experiments provide insight into subcellular localization (16) and copy numbers of proteins of interest (17–19).

Here, we characterize LexA mobility, quantity and heterogeneity *in situ* in unstressed cells and upon ciprofloxacin treatment with three different antibiotic concentrations reaching from 0.15× minimal inhibitory concentration (MIC) to 6×MIC. We capture molecule-to-molecule, as well as cell-to-cell variations in living and fixed bacteria using single-molecule tracking together with photoactivated localization microscopy (smtPALM). We identify four diffusive sub-populations, assign molecular functions using mutant LexA proteins and measure their time evolution. Complementing these live cell data, we develop an optimized single-molecule counting technique in *E. coli.* This approach allows us to quantify absolute LexA numbers in single cells and define the predominant location of LexA proteins within the *E. coli* cell. Combining live and fixed cell data, we devise a kinetic model which shows that under continuous antibiotic stress, copy numbers of individual LexA populations reach a new, unexpectedly high equilibrium, even under the lowest antibiotic stress. Our model provides for the first time *in vivo* LexA synthesis, cleavage, degradation and binding rates that are partially dependent on the antibiotic dosage. Finally, single-cell analysis reveals a homogeneous initial response to ciprofloxacin, and increased heterogeneity during late SOS response. We anticipate that this data and further experiments will help to shed more light on the bacterial SOS response and strategies to understand the development of antibiotic resistances.

## Results

### LexA assumes four diffusive states in *E. coli*

SOS gene transcription requires unbinding of the transcriptional repressor LexA. It is known that upon presence of single-stranded DNA, a RecA filament forms and triggers autocatalytic proteolysis of LexA and, as a result, a reduced occupancy of LexA at its transcription repression sites (Fig. 1a). Yet, little is known about the response kinetics of the LexA-SOS system to stress, in particular to antibiotics. To observe the bacterial SOS response in real-time, we developed an *in vivo* single-molecule reporter system on the mobility of LexA. We hypothesized at least two distinct LexA mobilities, a target site bound state and a freely diffusing state.

To test this hypothesis, we replaced the chromosomal *lexA* gene with a C-terminal *lexA-PAmCherry* fusion in *E. coli* MG1655. A structural analysis of LexA bound with its N-terminal domain to DNA (20) suggested that LexA function would be minimally disturbed by extending the C-terminal domain with a short six amino acid linker sequence prior to a photoactivatable fluorescent tag (Fig. 1b). The resulting strain showed normal growth, formed elongated filamentous cells under SOS response of equal dimensions as the wildtype and retained its susceptibility to ciprofloxacin (CIP), indicating a functional LexA-PAmCherry fusion that still allows repressor dimerization (Fig. S1a-d). Using a weak photoactivation with 405 nm illumination, a small number of LexA-PAmCherry (termed LexA^wt^ hereafter) repressors were fluorescent and single-molecule tracking was possible with an exposure time of 20 ms (Fig. 1c). Since clustering artifacts are sometimes observed with FPs of the mCherry family (19), we confirmed similar localization patterns of LexA-Dendra2 (Fig. S1e), an FP tag that is reported to be strictly monomeric (21). In unstressed *E. coli*, we then localized (22) individual molecules and tracked (23) their movements until photobleaching occurred (Fig. 1c).

For each molecule, we calculated the distance traveled between consecutive frames and generated a probability distribution of mean distances per frame per track to observe the extent of diffusive variety (Fig. 1d, Fig. S2). Surprisingly, three distinct peaks were observed in unstressed cells with inter-frame distances of approximately 55, 100 and 180 nm. Additionally, a broad tail at larger distances per frame appeared, indicating potentially four mobility states of LexA in unstressed cells. While this observation indicates a rich diffusive landscape of LexA, the actual area explored by the molecules cannot be readily quantified from mean distance traveled distributions. We therefore calculated apparent diffusion coefficients (*D*^∗^) for each trajectory from the change in mean squared displacement over time (see *Methods*, Fig. S2). The resulting distributions of apparent diffusion coefficients spanned four orders of magnitude (Fig. 2a, Fig. S3a). We fitted logarithmic gamma function-based models with varying numbers of populations using the *D*^∗^ probability distribution (see *Methods*) and revealed with the Bayesian information criterion (BIC) an optimum of four states (Fig. 2a). When we stressed the cells with the antibiotic ciprofloxacin (CIP), which is known to activate the SOS response (24– 26), the fastest population was strongly increased suggesting that this population represents cleaved repressors. Interestingly, the mean distance traveled per frame, as well as the apparent diffusion coefficients, showed only negligible shifts between all experiments, even during antibiotic stress (Fig. S3a). Therefore, we pooled all data sets and determined four global apparent diffusion coefficients, 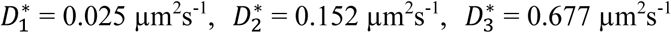 and 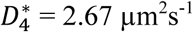 (Fig. 2a). We further applied an independent method to estimate the number of populations and their *D*^∗^ values. The user unbiased, Bayesian inference based algorithm SMAUG (27) collapsed from initially ten populations to a four-population model for both unstressed and stressed cells (Fig. 2c, Fig. S3b). The resulting diffusion coefficients agreed well with the logarithmic gamma functions fit to the diffusion coefficient histograms, yet it remained unclear what biological roles the four diffusive states represent.

**Figure 2:**
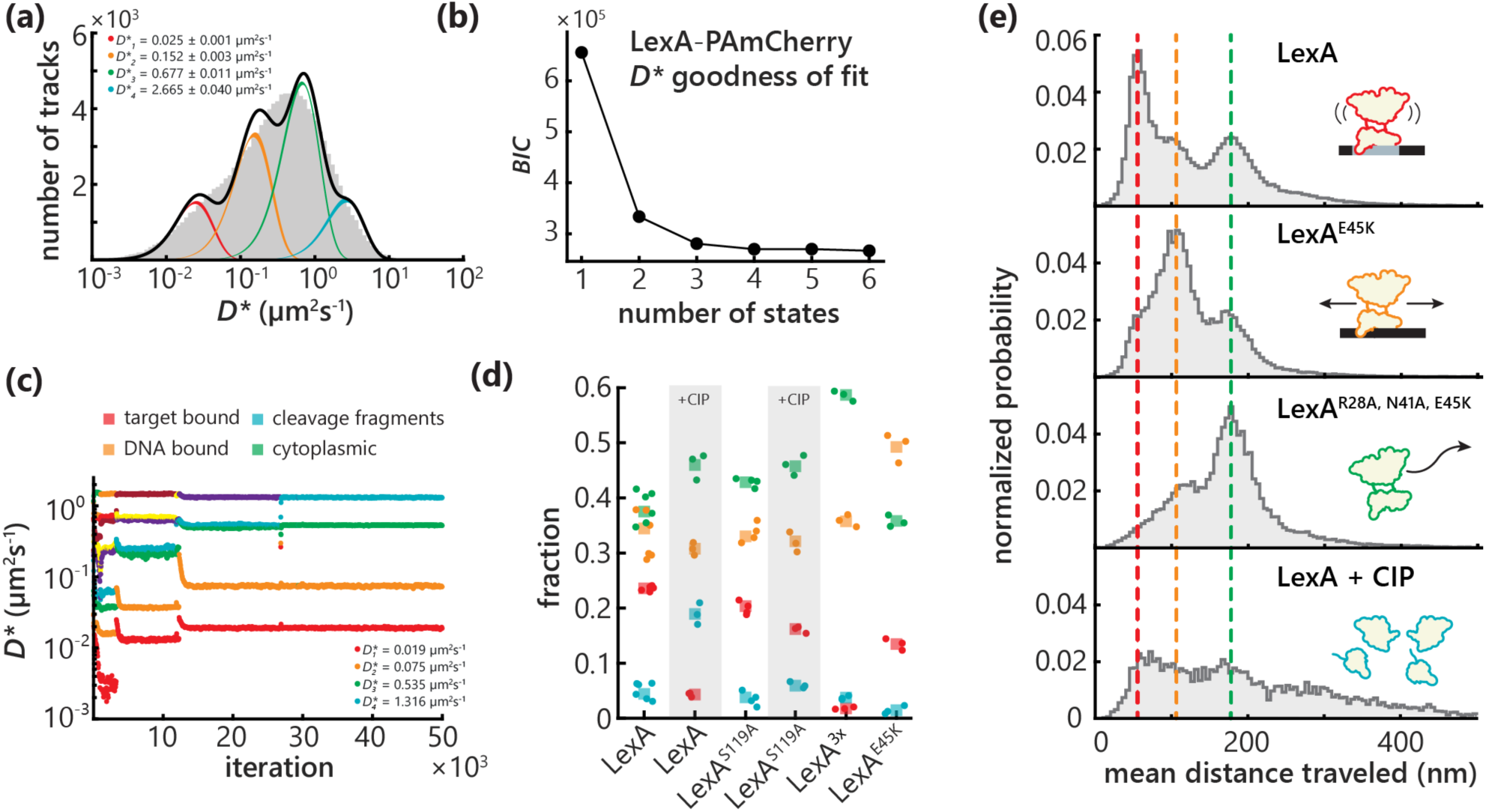
Identification of four diffusive LexA populations. **(a)** Determining population-specific diffusion coefficients by fitting a four-state model to pooled time course data (n = 401,875 tracks). Standard deviation is derived by bootstrapping. **(b)** Bayesian information criterion (BIC) for MLE fits of diffusive models assuming a varying number of states to *D*\* histograms. **(c)** Unbiased Bayesian analysis of LexA diffusion in unstressed cells using SMAUG (n = 73,580 tracks). Diffusion coefficients collapse to a four-state model after ∼26,000 iterations. **(d)** Contributions of the four populations in LexA mutants determined by MLE fits with fixed *D*\* values from (a). Color-coding according to (c) and (e). Squares represent the result of the global fit to pooled biological replicates (dots). Gray bars indicate addition of 20 ng/ml CIP for 45 min. **(e)** Histograms of mean distance traveled per frame by LexA^wt^ and mutants. Three peaks appearing in all mutants are marked by dashed lines. Pictograms indicate the proposed biological function for each dominant peak. The fourth, less sharp population increased upon 20 ng/ml CIP treatment for 45 min.

### LexA mutants reveal identities of four diffusive states

We devised a mutation-based strategy to identify the nature of the four observed states. We designed three LexA mutants containing the PAmCherry fusion: (i) LexA^E45K^, which was reported to abolish specific binding to the *recA* SOS box (28) and most likely other LexA operators but retains non-specific DNA binding; (ii) LexA^R28A, N41A, E45K^ (LexA^3x^), a previously unreported mutant, which should have a strongly reduced DNA affinity since two key residues involved in specific interactions with nucleobases (N41, E45) and one residue involved in backbone interaction (R28) were replaced (20); (iii) and LexA^S119A^, which is known to have an inactive catalytic center that renders the protein non-cleavable (29). We suspected the first two mutants to be lethal for *E. coli* due to their direct interference in the regulatory suppression of SOS genes. Therefore, we expressed LexA^E45K^ and LexA^3x^ from *P*_*lac*_ at the *lacZ* locus in addition to the untagged chromosomal *lexA*. The S119A mutation was inserted at the native locus on the chromosome (30) to serve as a negative control for SOS induction. We imaged all mutants under the same conditions as LexA^wt^ in unstressed cells, obtained localizations and tracked single molecules. We subsequently plotted the mean distance traveled per frame for all three mutants and used these distributions to assign biological functions to each population (Fig. 2e).

LexA^E45K^ shows a dominant peak at 100 nm mean distance traveled per frame, which corresponds very well to the second population of LexA^wt^. Knowing that LexA^E45K^ binds to DNA but weakly to target sites compared to LexA^wt^ (28), we infer that the second peak arises from DNA-associated LexA molecules, *i.e.* molecules that undergo a diffusional search along the chromosome to find their target site. The strong reduction in the 55 nm peak in this mutant suggests that the 55 nm peak represents target-bound LexA. The LexA^3x^ mutant showed a dominant peak at 180 nm mean distance per frame, coinciding perfectly with the third peak in the LexA^wt^ population. Considering that DNA binding is largely abolished in LexA^3x^, we assign this mean frame-to-frame distance to freely diffusing LexA dimers unassociated with the nucleoid. To identify the fourth and fastest population, which is underrepresented in unstressed cells, we treated *E. coli* with 20 ng/ml CIP for 45 min to induce a strong SOS response. Although this last population has no sharp peak in the mean distance traveled histograms, it is increasing and further expanding to larger mean distances upon stress, whereas the other populations diminish. We infer this population to be cleavage fragments (LexA^wt^ C-terminal domain fragments).

Next, we analyzed the LexA *D*^∗^ distributions of all mutant strains and quantified the relative abundance of each of the four diffusive populations and plotted their fractions (Fig. 2d). Upon antibiotic stress with 20 ng/ml CIP for 45 min, there is a strong increase in the fastest population. This population has a major contribution only in stressed cells, as expected for the increased concentration of cleavage fragments during SOS response. Target bound molecules, on the other hand, disappear almost entirely during stress, which is largely expected since unbinding of LexA is the major determinant of SOS gene expression. In contrast, the SOS response deficient LexA^S119A^ displayed only a minor difference between stressed and unstressed conditions. In the LexA^3x^ mutant, the target bound fraction dropped to the levels of SOS induction as expected. LexA^E45K^ showed only a small decrease in this population, most likely due to its residual affinity to stronger SOS boxes (28). It should be noted that, although the mutants are expressed from *P*_*lac*_ at likely much higher levels than LexA^wt^ from its native locus, heterodimer formation of untagged chromosomal LexA and the LexA^3x^ and LexA^E45K^ mutants cannot be excluded and might contribute to the target bound and DNA-bound populations. Overall, these results are in line with the qualitative observations based on the mean distance traveled histograms (Fig. 2e) and fit expectations for the respective biological function. In summary, we have identified four diffusive modes of LexA in live *E. coli* cells: target bound 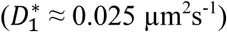, non-specifically DNA bound and most likely target searching 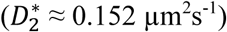, cytoplasmic diffusing and not DNA associated 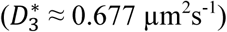, and cleaved LexA fragments 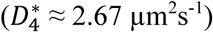.

### Dosage-dependent change in LexA dynamics upon ciprofloxacin treatment

In order to understand the dynamic regulation of LexA, we investigated the time-dependent change of each individual population during SOS response of different strengths. We determined the minimal inhibitory CIP concentration under our growth conditions to be ∼3 ng/ml (Fig. S1c), similar to previously reported values (31, 32). We restricted the treatment of *E. coli* with CIP to the exponential growth phase, since stationary phase cultures are less susceptible (33, 34). Cells were stressed with three different concentrations, 0.5, 3.0 and 20 ng/ml, *i.e.* ∼0.15×, 1×, 6× the MIC and imaged at eight incremental time points, 0, 10, 25, 45, 60, 90, 120 and 180 min for approximately 3.5 min. At least three independent biological replicates per time point and CIP concentration were recorded. For each experiment, we determined the *D*^∗^ of LexA^wt^ and the corresponding proportions of molecules in each of the four states. Corresponding histograms and fits for representative time points are shown in Fig. 3a for 20 ng/ml. We then plotted the fractions of each population over time for the three tested CIP concentrations (Fig. 3b-e). The variance between the biological replicates (dots in Fig. 3b-e) is small, illustrating high reproducibility. For all CIP concentrations, the target bound LexA^wt^ fraction declined rapidly during the first 45 min of stress, suggesting a fast response to DNA damage (Fig. 3b). DNA associated LexA^wt^ dimers declined as well within the first 45 min, but not as drastically (Fig. 3c). In contrast, the LexA^wt^ population of free dimers increased over time under all conditions (Fig. 3d). Cleavage fragments also increased rapidly and with a steeper slope at increasing CIP concentrations. However, since absolute LexA numbers change drastically during SOS response (see single-molecule counting below, Fig. 4c), kinetic rates such as cleavage and binding cannot be derived from relative population proportions only.

**Figure 3:**
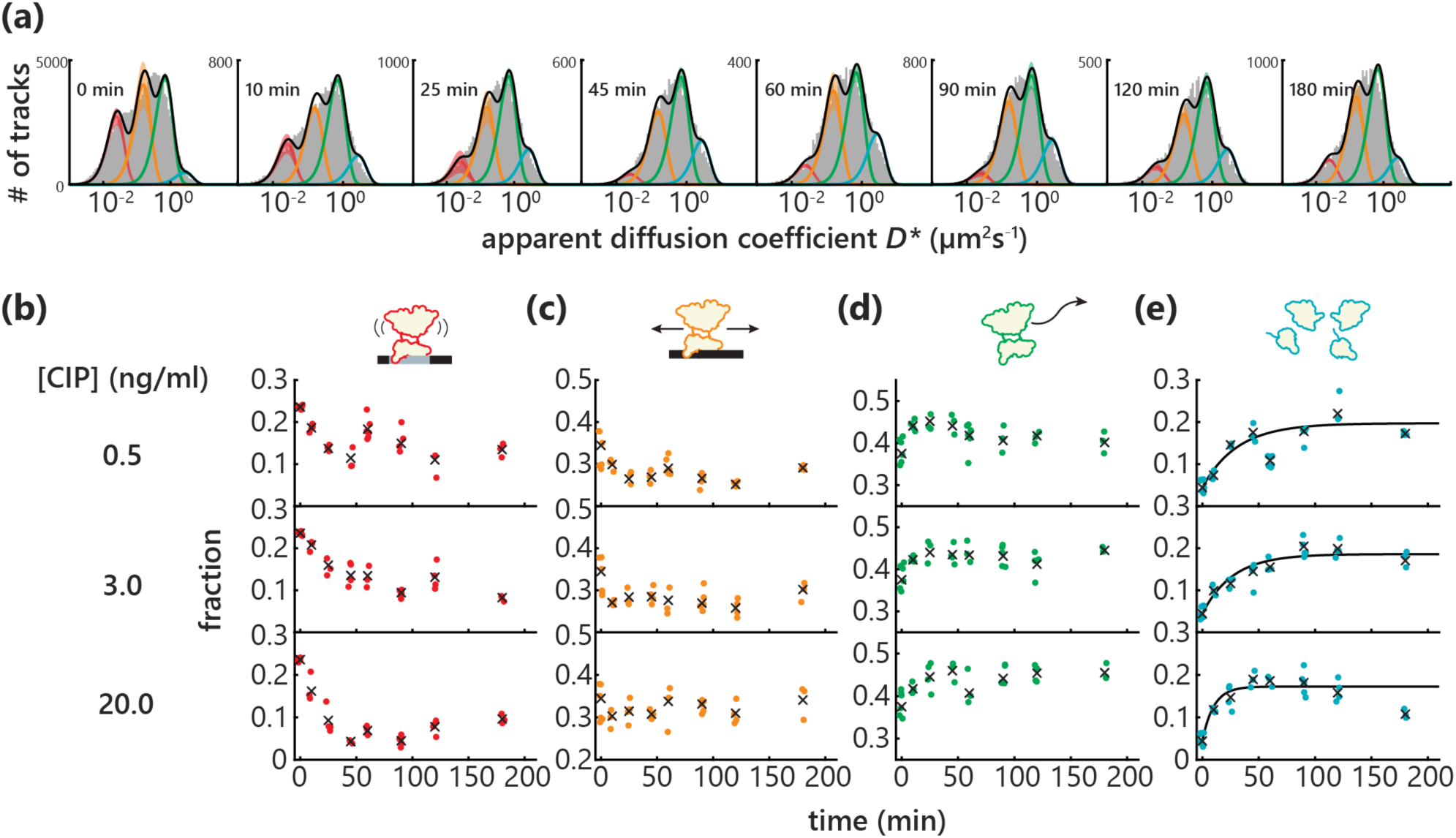
Temporal behavior of LexA diffusive populations during CIP stress. **(a)** Diffusion coefficient histograms of LexA during 20 ng/ml CIP stress, fitted with a global four-state model. Errors are standard deviations derived from at least three biological replicates. **(b-e)** Time course of contributions of each LexA diffusive population. Circles represent fits to individual biological replicates, crosses represent global fits to pooled replicates. Colors are according to pictograms above and to populations in (a). Black solid lines show exponential fits approximating *p*_*f*_(*t*).

**Figure 4:**
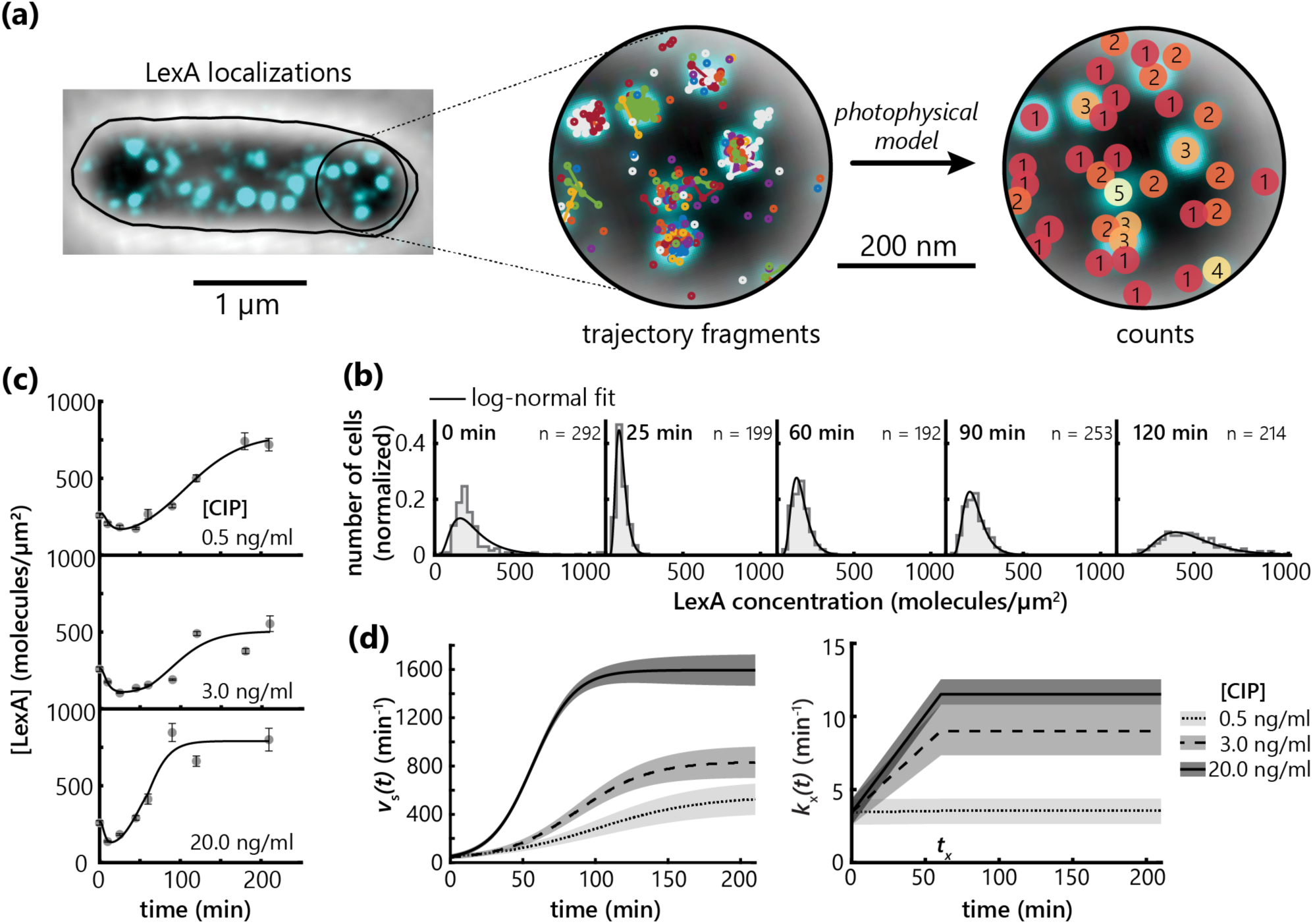
Single-molecule counting of LexA in fixed *E. coli*. **(a)** Localizations within a given radius are first connected in time to obtain trajectory fragments. The photophysical model with experimentally determined parameters is then applied to yield accurate molecular counts. **(b)** Histograms of cellular LexA counts normalized by cell area for 3 ng/ml CIP. Histograms are fitted with log-normal distributions to obtain statistic parameters. Counts are corrected for the detectability of PAmCherry in *E. coli*. **(c)** Time course of population mean for LexA concentrations obtained from log-normal fits as shown in (b). Error bars represent standard deviations of log-normal fits from 50 bootstrapping iterations. Solid black lines show the kinetic model describing LexA behavior under stress. **(d)** LexA synthesis and degradation derived from a simple model under indicated CIP concentrations.

### Single-molecule counting of LexA during SOS response

To obtain accurate single-cell LexA copy numbers during antibiotic challenge, and to obtain a full kinetic description of SOS repression, we applied single-molecule counting to quantify LexA^wt^. In fixed *E. coli* cells, we imaged LexA^wt^ using PALM until most FPs were bleached and only sparse photoactivation (approximately one localization per cell in 20 frames) was visible. After localization, we employed a counting strategy that is based on two previous approaches. First, an iterative algorithm (17) was used to extract the number of molecules within a small radius (Fig. 4a). This procedure requires knowledge of the photophysical parameters of PAmCherry, which were unknown for *in situ* experiments. We therefore applied a model of PAmCherry photophysics from activation, blinking and bleaching kinetics (Fig. S4a) inside fixed *E. coli* cells and determined all necessary parameters directly under our experimental conditions (Tab. S1, Fig. S4b-d). With these parameters at hand, we obtained spatially resolved fluorescence time traces (23) and predicted the number of single LexA proteins within one time trace based on iterative optimal-*τ*_c_ molecular counting as described by Lee *et al.* (17) (Fig. S4e, see also *Methods*).

All methods to determine absolute protein counts using only light emitting labels are biased by the detection efficiency of the respective fluorescent protein or dye (35). Improper folding, premature entry into the dark state and out-of-focus FPs or dyes can lead to a significantly large undetectable fraction. Knowledge of the stoichiometry of a labeled protein complex can be harnessed to determine this fraction (18, 36, 37). We introduced a known stoichiometry into the experimental system by labeling the well-characterized, strictly monomeric (38) inner membrane protein LacY with one or two copies of PAmCherry in tandem (tdPAmCherry). Both constructs showed clear membrane localization and LacY-tdPAmCherry portrayed significantly more double events within a single spot than LacY-PAmCherry, as determined with our counting algorithm (Fig. S4f). Using a global binomial fit, we determined a detection efficiency of 47.9 ± 6.6 %, which is much higher than previously shown for PAmCherry in *E. coli* (19), but in the same range as reported in *Xenopus* oocytes (36). We therefore corrected apparent single-cell counts by a factor of 2.09.

We applied this improved single-molecule counting method in fixed cells which grew under the same conditions as previously described for the live cell experiments. Raw LexA copy numbers per individual cell first decreased quickly, but then strongly increased during antibiotic stress (Fig. S3d). Before any CIP stress was applied, we detected 916 LexA molecules per cell (Fig. S3d), slightly lower than early estimates for *E. coli* MG1655 (6). This number converts to 458 dimers and approximately 108 target bound repressors using our estimated fraction for this population (23.6 %, Fig. 2d). This is in agreement with 57-102 reported LexA binding sites in the *E. coli* genome (3).

Previous studies observed that the SOS response leads to a phase of nucleoid compaction and replication inhibition (39), followed by DNA dispersion (40) and possibly resumption of replication. Filamentous *E. coli* therefore may contain multiple chromosomes (41) and the cytoplasmic content of several normal-sized cells. To account for this possibility, we decoupled LexA copy numbers from cell size and cycle by normalizing each count with cellular area derived from brightfield images. Distributions of LexA concentrations remained unimodal throughout the SOS response and could be approximated with a log-normal probability density function (Fig. 4b, Fig. 6b). This allowed us to determine the expected value, which we used as the number of LexA proteins per cell area for each condition (Fig. 4c). We observed a fast, CIP concentration-dependent degradation of LexA^wt^ within the first 25 min, where minimal levels of 100-150 molecules per µm^2^ were reached (depending on antibiotic dosage), corresponding to ∼300-500 molecules per cell on average. Given that at least 57 SOS genes require at least two LexA monomers for repression, such low numbers cannot sustain SOS repression. Accordingly, rapid degradation is followed by strong new synthesis due to net dissociation of the repressor from its own operator and subsequent *lexA* expression. This fast initial degradation also underscores that the LexA C-terminal cleavage fragment, which remains fused to PAmCherry after auto-proteolytic cleavage, is efficiently degraded by the corresponding proteases. At 20 ng/ml CIP, the concentration of LexA^wt^ molecules per cell increased after 200 min incubation to 4-fold of the unstressed level (Fig. 4c). Interestingly, a strong increase was also observed at 0.5 ng/ml CIP (0.15×MIC), indicating a potent SOS response even at low continuous antibiotic dosage.

LexA^wt^ counts over time contain both degradation and synthesis of new LexA^wt^ and can be modeled using a simple three-state reaction scheme with irreversible steps. Central is the visible state *V*, where the molecules are detectable using fixed-cell PALM (Fig. 5a, see also *Methods*). The influx to this state *v*_*s*_ represents the new synthesis of LexA^wt^ while the outflux 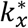 represents the apparent degradation rate of all LexA states together (*V*). From the time evolution of our first set of experiments in live *E. coli*, we extracted the LexA^wt^ cleavage fragment probability function *p*_1_(*t*) (Fig. 3e) and could therefore determine the degradation rate of LexA^wt^ fragments *k*_*x*_ (see *Methods*). We tested different response functions for *v*_*s,CIP*_(*t*) and found that a simple logistic function (*Methods*, Eq. 3) describes the delayed new synthesis well (Fig. 4c and d, left). Increasing CIP stress lead to a faster response time (Fig. 5g) and stronger increase in the synthesis rate followed by a higher synthesis saturation speed, creating independent new equilibria of the regulatory network. For the degradation rate *k*_*x*_, we initially assumed a time- and stress-independent degradation rate, which could not describe our data well (Fig. S6a). By introducing a stress-dependent response function 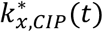 (*Methods*, Eq. 1iii) for the degradation rate, a better description of the experimental data with minimal number of parameters could be achieved. Since there is insufficient information available for the regulation of *E. coli* degradation complexes, we assumed a simple linear increase of the degradation rate followed by a stress-dependent saturation at time *t*_*x*_ (Fig. 4d, Fig. S5b). The degradation speed *v*_*x*_(*t*) = *p*_*F*_(*t*) · *k*_*x*_(*t*) · [*V*] at the new equilibrium agrees with the synthesis of LexA^wt^ monomers at ∼1600/min at 20 ng/ml CIP. We can conclude that the SOS response reacts to different external stress levels within 100 min (Fig. 5g) and returns the system to equilibrium within 110-150 min, even when the stress level exceeds the MIC 6-fold and the cells will likely not recover. Having this quantitative data at hand, we proceeded to dissect the regulatory feedback further using the four diffusive states of LexA^wt^.

**Figure 5:**
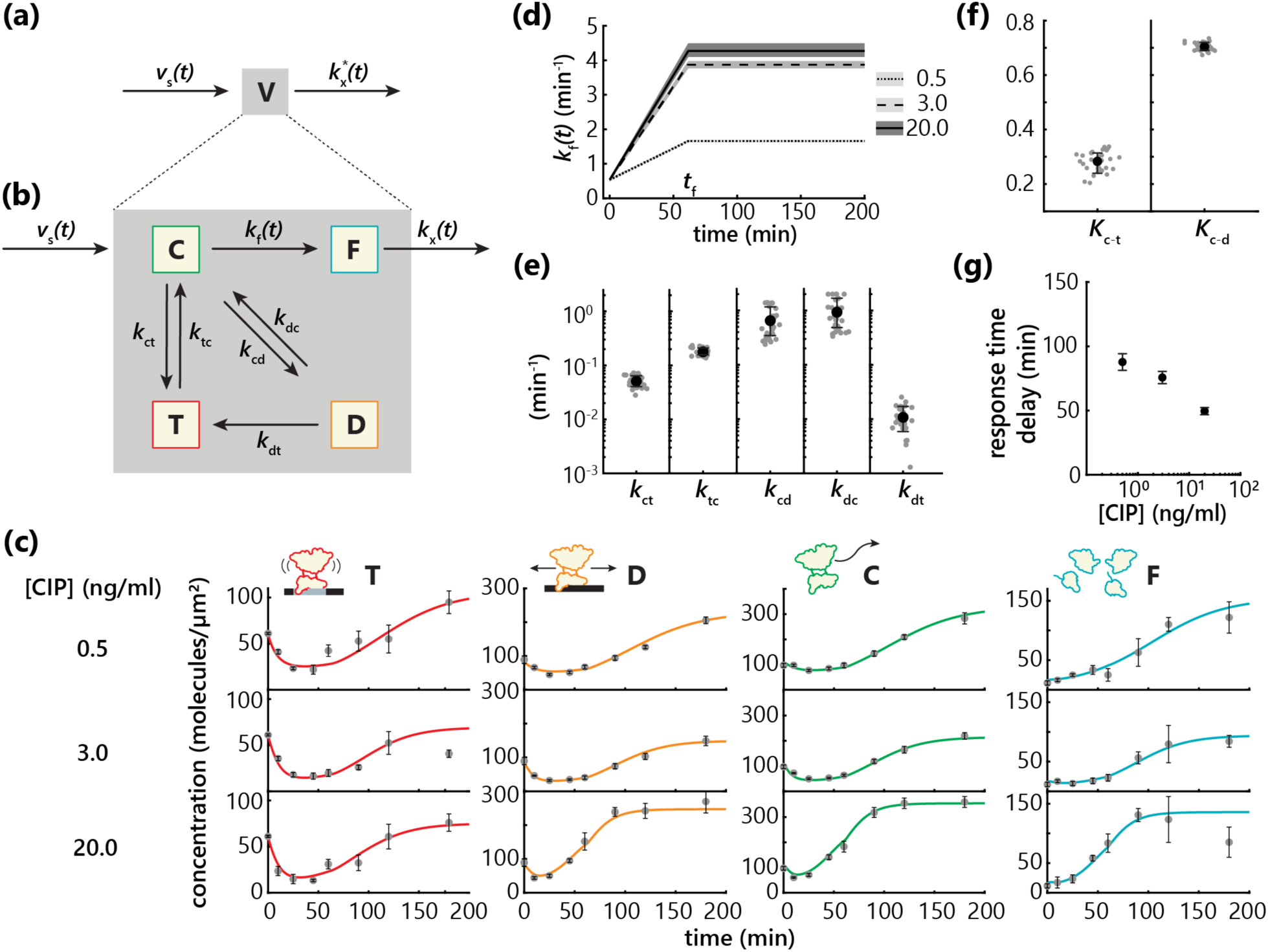
A kinetic model reveals transition rates in a network of LexA diffusive states. Model of LexA synthesis and degradation to describe single-molecule counting data. **(b)** Full model to describe the progression of each individual LexA population throughout the SOS response. T: target bound, D: DNA bound, C: cytoplasmic, C: cleavage fragments. **(c)** Concentrations of individual populations over time at indicated CIP concentrations. Solid lines represent the fit of the kinetic model shown in (b). **(d)** Change of the cleavage rage *k*_f_ over time, modeled with a step function. **(e)** Individual rates according to (b). Errorbars represent standard deviations from 30 bootstrapping iterations using different combinations of biological replicates. **(f)** Equilibrium constants for LexA DNA binding reactions. **(g)** Response delay between decay of target bound LexA and synthesis of new molecules dependent on CIP concentration.

**Figure 6:**
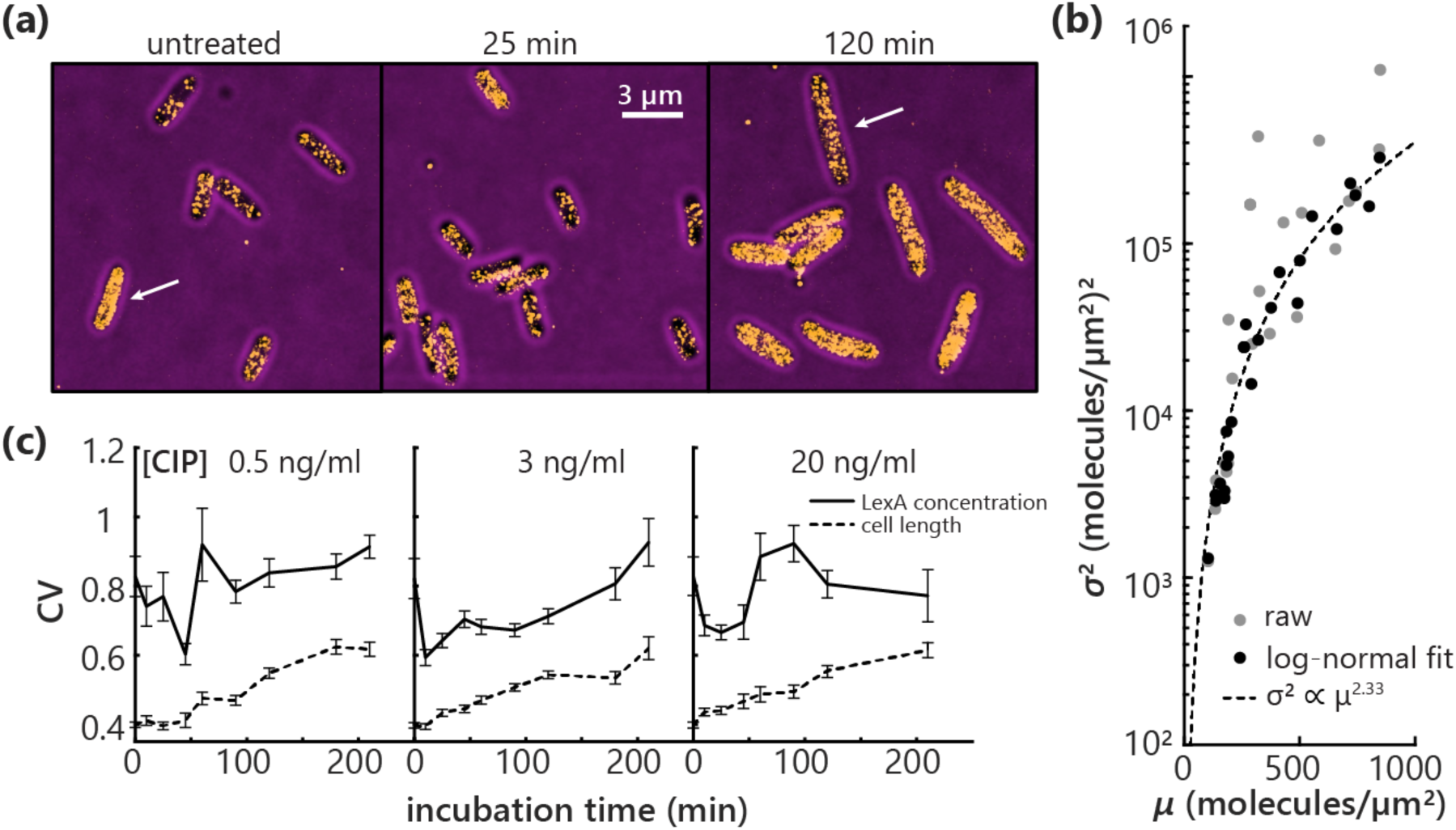
Cell-to-cell heterogeneity during SOS response. **(a)** LexA PALM images in fixed cells, superimposed on brightfield images at 20 ng/ml CIP treatment for indicated incubation times. Arrows highlight cells with specifically high or low LexA concentrations. Bright spots outside of cells are fiducial markers used to correct for sample drift. **(b)** Relationship of variance (*σ*^2^) and mean (*µ*) derived from LexA concentration distributions, either raw or from a log-normal distribution fit. Each point represents one experimental condition, *i.e.* varying time and CIP concentration. The dashed line shows a power-law fit to the parameters derived from log-normal fits. **(c)** Coefficient of variation (CV) for cellular LexA concentrations and cell length derived from log-normal fits. Errorbars represent standard deviations derived from 50 bootstrapping iterations.

### A kinetic model of transitions between LexA diffusive states

To determine *in vivo* interconversion rates between the four diffusive states, we expanded the coarse-grained state of visible molecules *V* to four diffusive states: target-bound (T), DNA-bound (D), cytoplasmic and LexA^wt^ C-terminal fragments (F) (Fig. 5b). The single-molecule counting data allowed us to convert the time evolution of probabilities into molecular densities (LexA^wt^ per µm^2^) for each CIP concentration (Fig. 5c). In our initial assumption, we coupled all four states with each other through differential equations. Based on earlier reports (8), the cleavage reaction to LexA^wt^ fragments is only possible from cytoplasmic LexA^wt^ and irreversible, therefore we could simplify the set of equations (*Methods*, Eq. 6). We then assumed reversible transitions between the remaining three states, C, T and D. All transitions between C, T, D, and F depend on the population of the outgoing state, except that C will be provided with the influx from new synthesis and F will be depleted by degradation. Both, synthesis and degradation were determined by the fit result of the counting data in the previous section. Using global optimization considering all tracking data sets at the same time and deriving errors by drawing iteratively from the pool of biological replicates, optimal rates for each transition were determined using a reduced 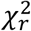 criterion for the fit quality. In order to reproduce our data, the transition rate *k*_1_ needed to be time-dependent, which is plausible because CIP stress generates time-dependent DNA damage and ssDNA exposure, leading to time-dependent auto-proteolytic cleavage. We modeled the increasing fragmentation rate *k*_1_(*t*) similar to *k*_*x*_(*t*) with a ramp function (*Methods*, Eq. 5iii, Fig. 5d, Fig. S6b). Using this model adjustment, we could reproduce in a global approach for each CIP concentration the LexA^wt^ densities of the four states (Fig. 5c). The different transition rates between C, T and D covered about two orders of magnitude (Fig. 5e), while each replicate condition produced a similar set of rates (gray dots in Fig. 5e), suggesting high reproducibility. Surprisingly, our model revealed a low transition rate from D to T, which could be reasoned with current models of target search in the genome, that a mere 1D search in DNA is unlikely to succeed in a rapid target binding (42). This is further supported by the relatively high transition rate from C to T, which we initially assumed to be low considering the number of LexA binding sites and the genome size of *E. coli.* Furthermore, the transition rate from T to D appeared to be neglectable, which might indicate that the direct dissociation to the cytoplasm is preferred over the back transition to the DNA-bound state. Overall, the equilibrium between the cytoplasmic and the target bound state *K*_*C*−*t*_ was ≈0.28, indicating a bias towards a cytoplasmic population and regular exchange of transcriptional repressors (Fig. 5f). Noteworthy, our model assumed that the transitions between C, D and T are independent of CIP concentration, and that CIP stress affects only the synthesis speed *v*_*s,CIP*_(*t*), the fragmentation rate *k*_*f,CIP*_(*t*) and the degradation rate *k*_*x,CIP*_(*t*). The combination of these together with the binding and unbinding kinetics of LexA^wt^ to DNA and its target site lead to differential response times after CIP exposure in *E. coli.* While the response time delay, the time lag between initial LexA cleavage and new synthesis, is ∼80 min at sub-MIC (0.5 ng/ml CIP) stress, the cells responded after ∼40 min at 6×MIC (20.0 ng/ml) CIP under slow growth conditions in minimal medium.

### Cell-to-cell heterogeneity during SOS response

Within the SOS network, heterogeneity in repressor copy numbers might arise from several processes including stochasticity or availability in RecA filamentation and subsequent LexA proteolysis, on-off binding rates of the LexA repressor on its own promoter (9) or transcriptional bursting during *lexA* expression itself (43). Time-resolved single-cell repressor counts and concentrations revealed varying amounts of heterogeneity throughout the SOS response (Fig. 4b, Fig. S3d). Notably, varying concentrations between individual cells were visually discernible in PALM images of fixed cells (Fig. 6a).

For *E. coli*, the scaling of protein density variance *σ*^2^ with mean protein density *μ* has previously been described with the power law *σ*^2^ ∝ *μ*^1.5^ (44, 45). We observed that during the SOS response *σ*^2^ of [LexA] follows a power-law scaling, but with a larger exponent of 2.33 indicating unusually large heterogeneity (Fig. 6b). This is likely due to the self-regulatory feedback of LexA. Our data also allowed us to quantify the time-resolved heterogeneity by calculating the coefficient of variation *CV* from log-normal fits to 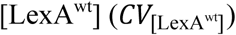) and cell length distributions (*CV*_length_), using 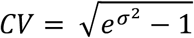 with *σ*^2^ corresponding to the variance of the log-normal distribution. The *CV* yielded a relative measure of variance at each time point of CIP stress (Fig. 6c). The cell length decreased initially after 10 min of CIP treatment at all concentrations (Fig. S1f), likely due to rapid arrest of cell division, however, the *CV*_length_ monotonically increased over time for all [CIP] conditions, independently of CIP concentration. In contrast, the heterogeneity of [LexA^wt^] showed a strong drop within the first hour of SOS response compared to the steady state of unstressed cells under all antibiotic concentrations. In the second hour of incubation, 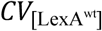 displayed a brief peak and at longer CIP incubation times 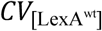 stabilized approximately at the unstressed level, indicating a new stable feedback. Since the first hour is dominated by LexA loss rather than synthesis (Fig. 4c), the decrease in cell-to-cell variation within this period and the subsequent increase with more repressors being synthesized (Fig. 4d) indicate LexA unbinding and synthesis as the major sources of variability, rather than signaling events upstream.

## Discussion

The underlying mechanism of SOS response activation (2, 3), as well as the gene expression profile of SOS activated cells (46) have been extensively studied. Regulatory steps leading to these patterns, kinetics of regulation in living cells, as well as cell-to-cell variation of SOS induction are still lacking a global understanding. Here, we studied the dynamics of the master-regulator of the SOS response, LexA, in non-stressed cells as well as during continuous DNA damage with the antibiotic ciprofloxacin. We used single-molecule tracking in live cells, single-molecule counting in fixed cells and a set of mutants to characterize the stress response under low, medium and high antibiotic stress. We present here the course of SOS response activation and feedback regulation on the level of binding, unbinding, cleavage and synthesis of the master repressor LexA itself.

Four diffusive states, even in unstressed cells, represent a surprising variety in the LexA mobility landscape. By comparing several mutants of the repressor in their diffusive pattern, we identified the four states as target bound, non-specifically DNA bound, cytoplasmic, and cleaved LexA fragments. The non-specifically DNA bound state is supported by previous reports that prior to finding the target sequence and firmly binding to it, repressors often associate with the DNA in a sequence non-specific manner and perform a 1D target search, as previously shown for the *lac* repressor (47, 48). Tight binding to operator sites results in an apparent diffusion coefficient 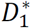 close to zero, defined by minimal movement of DNA within bacterial cells (49, 50). The number of tightly bound repressors quickly decreased already after 10 min of CIP treatment (Fig. 5c), showcasing the prompt response to antibiotic stress and DNA damage under exponentially growing conditions. Our quantitative model suggests that every ∼6 min, LexA is unbinding the target bound state to the cytoplasm, which allows for the short reaction time. This rapid regulation was previously difficult to capture on the cellular level due to the time delay inherent to indirect fluorescent reporters, though observed on the mRNA level for several SOS genes (46), well in line with our finding. The target-bound population decreased more rapidly when the bacteria were exposed to higher concentrations of CIP (Fig. 5c). This dose-response relationship of the SOS response has been previously observed for pulses of UV damage (6, 9), but is not well described for continuous antibiotics exposure. Counting of LexA levels during CIP stress agreed qualitatively with previously reported pulsed UV exposure stress (6, 51–53). Surprisingly, even in unstressed cells, we found about 4 % of LexA in the cleaved fragment population, suggesting a low level of LexA fragmentation under normal growth conditions with a small, but detectable LexA turnover.

Both the target bound and the non-specifically DNA bound state quickly diminish during CIP exposure (Fig. 5c), as overall cellular LexA concentration drops and cleavage and degradation dominate during the early SOS response, especially at above-MIC antibiotic concentrations. Interestingly, we observe that particularly the target bound LexA population recovers during late SOS response, as an immense amount of LexA up to 800 molecules per µm^2^ is synthesized and cytoplasmic concentrations increase. A new equilibrium is reached after 100-120 min, depending on CIP concentration, as LexA synthesis balances continuous cleavage. It follows that *E. coli* returns to repressing SOS genes before intense mutagenesis and cell-cycle inhibition lead to death, assuming resistance-granting mutagenesis took place within the time frame of maximum upregulation. At sub-MIC CIP, this recovery is significantly slower, suggesting the bacterial attempt to carry out mutagenesis and adaptation in a more controlled fashion.

By applying a simple kinetic model (Fig. 5b) to this time-course data, we were able to determine the key parameters fully describing LexA action *in vivo*. The rates associated with operator binding represent mean rates over all LexA binding sites, which display large variations in binding affinity (9). They did not show variation under CIP stress, and we therefore propose that cleavage, degradation and synthesis rates are the major determinants of LexA abundance during SOS response. We determined a maximum cleavage rate *k*_1_ as 4.3 min^-1^ (Fig. 5d), well in line with maximum cleavage rates of 3.6-4.2 min^-1^ measured *in vitro* (54, 55). This rate is higher for higher CIP concentrations likely due to increased cytoplasmic LexA availability and formation of RecA* filaments with increasing DNA damage (Fig. 5d). Degradation of LexA fragments is mediated by the ClpXP protease complex (10). Our results suggest that this process must be highly efficient *in vivo*, given the high degradation rates (3.5-11.5 min^-1^) that explain the fast decay of total LexA in the early SOS response, as well as counter the accumulation of cleavage fragments (Fig. 4d). Averaged PALM images reveal localization of LexA towards the pole regions (*Supplementary Information*, Fig. S5), which is also where ClpXP resides during heat shock stress to dissolve protein aggregates that are occluded from the nucleoid (56). This localization might also stem from freshly synthesized repressors still in proximity of ribosomes (57). We note however that we found no evidence of LexA aggregates in single cells, and the average images (Fig. S5) merely present trends of preferential localization overall. Interestingly, the LexA^wt^ saturation rate for synthesis at 20 ng/ml CIP is ∼1,600 LexA^wt^ per minute for a 444 amino acid (aa) protein. Taking into account previous reports of translation rates of ∼15 aa/s (58), this would mean that ∼800 ribosomes are occupied with LexA^wt^ synthesis, which is 2 % of all available ribosomes in *E. coli* (57) to maintain the new equilibrium at high stress conditions.

Cell-to-cell heterogeneity is a major driver of microbial evolution and thought to play an important role in antibiotic resistance acquisition (59). Within the SOS response, heterogeneity has been observed before with fluorescent reporters expressed from the SOS promoters (13, 60). Reporting directly on single-cell protein levels with *lexA* expressed from the native genomic locus, we showed large LexA copy number heterogeneities, and thereby likely heterogeneity of downstream transcriptional output, which varies during antibiotic challenge (Fig. 6). An initial decrease of the *CV* highlights a uniform response to initial damage. Downstream events such as noisy gene expression from SOS promoters (12) likely lead to increased heterogeneity during the later response and with it, chances of population survival increase rather than those of a single cell. Our data further suggests that regular un- and rebinding of LexA occurs on the order of ∼10 min in unstressed cells, likely causing sporadic expression of SOS genes and might thereby increase mutagenesis events that lead to persistence (61, 62) even before antibiotic exposure.

In conclusion, we directly observed individual states of the SOS response master regulator over time with single-molecule tracking and quantitative PALM during constant ciprofloxacin stress. We discovered that the response kinetics of individual processes differ greatly with increasing stress, however, *E. coli* manages to re-establish a new equilibrium under all tested conditions within ∼100 min. We further provide regulatory timescales and precise LexA quantities that help to better understand the bacterial response to DNA damaging antibiotics and might contribute to developing new strategies for SOS response inhibition. In the future, our *in vivo* quantities can help to parametrize more complex simulations of the whole SOS network to finally achieve a systems-level understanding of this intriguing network. The combined approach of single-molecule tracking and counting can be adapted to a range of different proteins to monitor molecular sub-populations and corresponding expression changes in a precise manner.

## METHODS

### Bacterial strain construction

To tag all genes at their endogenous locus, the gene for PAmCherry was cloned into the plasmid pR6K-lox71-cm-lox66 by ligation into a unique BstAPI restriction site. For LexA mutants that were integrated at the *lacZ* locus, the *lexA* gene was amplified from the *E. coli* MG1655 genome by colony PCR and cloned into pR6K-PAmCherry-lox71-cm-lox66 using the In-Fusion^®^ HD cloning kit (Clontech). Mutations were introduced using the QuikChange II Site-Directed Mutagenesis Kit (Agilent). A targeting cassette including ∼50 bp homology to the target region on either end was amplified by PCR. We described the construction of the genomic LexA^S119A^ mutation previously (30). Recombineering (63) was carried out using a protocol adapted from (64). *E. coli* MG1655 was transformed with pSC101-BAD-γβαA, where Redγβα expression was induced with 0.25 % L-arabinose. 250 ng of the targeting cassette were electroporated into competent cells. The temperature-sensitive plasmid was removed by incubation at 37 °C. The genotype of successfully engineered clones was verified by colony PCR and sequencing. The chloramphenicol resistance gene was recycled using Cre recombination from the temperature-sensitive plasmid pSC101-BAD-Cre.

### Culture conditions and sample preparation

Bacterial cultures were grown in M9 minimal medium (1 x M9 salts [Sigma], 1 mM MgSO_4_, 0.1 mM CaCl_2_, 0.4 % glucose) supplemented with 0.001 % biotin and thiamine. Autoclaved 1.5 ml tubes with a punctured lid were used for incubation in an Eppendorf ThermoMixer^®^ at 37 °C and 950 rpm. For imaging, cultures were streaked on LB plates without antibiotics from a glycerol stock. 1 ml M9 cultures were inoculated from the plate and grown over night. The cultures were diluted 36-fold into 1.44 ml fresh M9 medium and grown until exponential phase for 3.5 h. If applicable, the lactose operon was induced with 150 µM IPTG at the beginning of incubation for imaging of LexA mutants at the *lacZ* locus or LacY-PAmCherry, and with 17 µM for LacY tandem constructs. Bacteria were then treated with CIP and incubation was continued as indicated.

For live-cell experiments, cultures were then centrifuged for 30 s at 7,300 g, most of the medium was removed and the pellet was resuspended in an appropriate amount of M9. 1.5 mm thick coverslips were cleaned at least twice by sonicating for 15 min in 5 % Mucasol, 15 min in ethanol and drying with nitrogen gas. An agar pad containing M9 medium as well as CIP, if applicable, was prepared according to (65). 2 µl of the concentrated cell suspension were placed on top of a clean coverslip, covered by the agarpad and immobilized by adding another coverslip on top and pressing gently.

For fixed-cell experiments, cultures were grown as described above, cooled on ice for 10 min and incubated in 4 % paraformaldehyde in PBS, pH 7 for 35 min at room temperature, then washed twice with PBS and stored at 4 °C. Prior to imaging, dark red Fluospheres 0.2 µm (Thermo Fisher) were diluted 2,000-fold into the concentrated cell suspension and the sample was assembled as described above.

### Data acquisition

PALM movies were acquired at room temperature on a Nikon Ti-E STORM/PALM microscope equipped with a 1.49 NA 100 x TIRF objective (Nikon) and Andor iXon Ultra 897 EMCCD camera. PAmCherry was continuously activated with a 405 nm laser and excited at 561 nm with 28 mW (measured after the objective) under near-TIRF conditions. The 405 nm activation laser was increased over the course of a movie following a Fermi function as described previously (17). Emission light was filtered by a 605/70 bandpass for fixed-cell movies or by a 575 longpass (AHF) for live-cell experiments to collect more photons from diffusing PAmCherry fusions. For single-molecule tracking experiments, around 10,000 frames were recorded at 20 ms exposure time. Fixed-cell PALM movie length varied according to the protein copy number between 20,000 and 30,000 frames. Brightfield images of the same field of view were taken after PALM acquisition.

### Raw data processing

Raw movies were processed using SMAP (22) with settings for 2D localization (parameters listed in Tab. S2). The shape of the canonical point spread function was calibrated from 20 z-stacks (10 nm step size) of dark red fluorescent beads under the exact experimental conditions. Cells were segmented from brightfield images using Oufti (66). Localizations were then grouped based on the *E. coli* outlines using custom-written MATLAB script.

### Live-cell data processing

The tracking software swift (23) was used with the parameters listed in Tab. S2 to obtain LexA-PAmCherry trajectories. Tracks with less than four and more than 50 localizations were discarded. To create histograms from the mean distance traveled per frame, we took the mean of all step lengths within a track and weighted the histogram according to the number of steps (Fig. S2). Apparent diffusion coefficients were calculated from track fragments of four localizations in length (Fig. S2). If a track was at least eight localizations long, a maximum of three fragments were recovered from a single track. Apparent diffusion coefficients (*D*^∗^) were extracted from mean squared displacement (*MSD*) over time lag (*τ*) curves using linear fitting according to

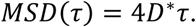

The *MSD* was calculated for each *τ* using the following equation, with *x*_*i*_ and *y*_*i*_ being coordinates of the consecutive localizations making up a trajectory.

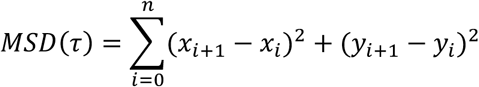

Distributions of *D*^∗^ were fitted with analytical equations as first described by Vrljic *et al.* (67), composed of *r* linear combinations of gamma-distributions:

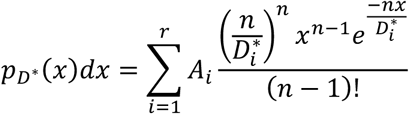

with *A*_*i*_ and 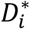 being the probability and average apparent diffusion constant of state *i*, respectively, and *n* being the number of steps, in this case three (four localizations). The fits were performed and displayed at logarithmic x-axis using a linear combination of *r* log-transformed gamma distributions.

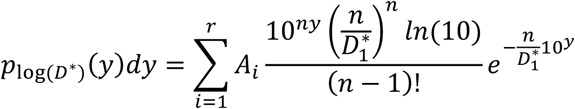

Maximum likelihood estimation (MLE) was realized in MATLAB and the Bayesian information criterium (*BIC*) was determined directly from the maximum value of the likelihood function 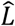 with *k* being the number of free parameters and *m* being the sample size.

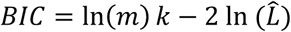

*D*^∗^ distributions from unstressed wildtype cells were fitted in this way to determine the optimal size of the analytical model based on a minimized *BIC*. A four-population model was found most probable, which was confirmed by applying SMAUG (27) to the same dataset (Fig. S2). The four apparent diffusion coefficients were determined from pooled *D*^∗^ values, and globally fitted with a four-population model. Standard deviations were determined from 50 bootstrapping iterations by randomly drawing *m* datapoints from each dataset with replacement; *m* is the size of the smallest dataset. The four *D*^∗^ parameters obtained in this way were fixed and only *A*_1_-*A*_3_ fitted to single biological replicates. Biological errors were based on standard deviations of at least three biological replicates. Additionally, data from all replicates within one dataset were pooled and fitted again to obtain more accurate, global amplitudes. The sample sizes together with mean distance traveled and *D*^∗^ distributions of all live-cell data collected are recorded in Fig S3.

### Fixed-cell data processing

For fixed-cell movies, sample drift was corrected using red-shifted fluorescent beads as suggested recently (68). A custom implementation of the “DriftCorrectionFiducial” plugin from PALMsiever (69) was employed to directly shift PAmCherry localizations based on smoothed frame-to-frame translation of the fiducial markers. Cells were manually sorted into dividing and non-dividing based on the presence of an indentation at the cell mid. Single-molecule localization images were rendered with the ThunderSTORM plugin for ImageJ (70).

#### Extraction of in situ photophysical parameters of PAmCherry

Fluorescence time series of active fixed PA-FPs were extracted from raw movies by swift (23) using the following set of parameters: *exp_displacement* = 55 nm, *max_displacement* = 125 nm, *max_blinking_duration* = 10 frames, *precision* = 55 nm, *exp_noise_rate* = 10 %, *p_switch* = 0, *p_bleach* = 0.0001 and *p_blink* = 0, where the latter two parameters were set to zero to avoid the connection of fluorescent traces interrupted by dark frames. Next, the lengths of all fluorescence segments were sorted in a histogram (Fig. S4b). The resulting *T*_on_ distribution is well described by the sum of two exponential functions (*R*^2^ = 0.99):

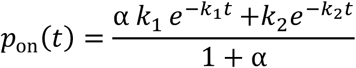

with *α* and 1/(1 + *α*) being the fractions of segments with rate *k*_1_ and *k*_2_, respectively. The corresponding average rate ⟨*k*⟩ is directly connected to the sum of the bleaching rate *k*_*b*_, and the transition rate from the active to the dark state *k*_*d*_. In order to obtain the recovery rate *k*_*r*_, from the dark to the active state, the spatial positions of the segments were correlated: all segments with a distance smaller than 125 nm were connected. From the resulting time traces a *T*_off_ histogram was collected from the length of fluorescence gaps (Fig. S4c). The resulting distribution contains a mixture of blinking events and activation of new PA-FPs within the same radius and is well described by a sum of three exponential functions (*R*^2^ = 0.96):

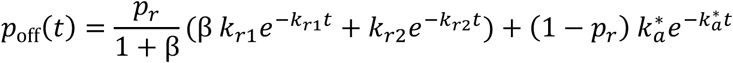

While the biggest two relaxation rates, *k*_*r*1_ and *k*_*r*2_ (>1 s^-1^) most likely correspond to the recovery of the PA-FPs from the dark to the active state, the smallest relaxation rate 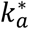 of about 0.1 s^-1^ refers to the activation of new fluorescence proteins. The ratio of the two species is given by *p*_*r*_. To account for the new activation of PA-FPs, a blinking tolerance time *τ*_*c*_ = 750 ms was chosen from the *T*_off_ distribution (Fig. S4c). Subsequently, the number of blinking events were collected from the separated fluorescence time traces and sorted in an *N*_blink_ histogram (Fig. S4d), which was then analyzed as described earlier (17). The obtained kinetic parameters are listed in Tab. S1. For further counting experiments, the reappearance probability *P*_reappear_ = Δ*t*/⟨*t*_off_⟩, the blinking probability *P*_blink_ = Δ*t*/⟨*t*_on_⟩ and the bleaching probability *P*_bleach_ = *k*_*b*_/Δ*t* were calculated and used as experimentally determined input parameters for swift.

#### Single-molecule counting

In order to quantify the number of molecules per localization spot of the recorded movie, first trajectories were built with swift using the parameters as in the previous section. From the localization list, an activation profile (number of active molecules per frame) was accumulated (Fig. S4e, top) and described using polynomial fits. Next, the calibration curve *τ*_*c,opt*_(*N*_mol_) was obtained from Monte Carlo simulations using the photophysical parameters of PAmCherry (Tab. S1) as previously described (17), where *N*_mol_ denotes the number of simulated molecules and *τ*_*c,opt*_ the optimal blinking tolerance time, where over- and undercounting cancel out each other. Briefly, in the Monte Carlo simulation, *N*_mol_ was varied from 2 to 100 molecules. For every iteration, *N*_mol_·5000 fluorescence traces were created by first drawing activation frames from the approximated activation profile (Fig. S4e, top). Subsequently, for every molecule a sequence of active and dark dwell times was drawn using the experimentally derived *p*_on_, *p*_off_ and *N*_blink_ distributions (Fig. S4b-d). The resulting *N*_mol_·5000 traces were randomly merged to simulate *N*_mol_·5000 molecules per localization spot. The optimal blinking tolerance time *τ*_*c,opt*_ was then chosen such that the average number of molecules per spot matched the simulated number of molecules (Fig. S4e, bottom). Revisiting the localization list, localizations with a distance smaller than 55 nm were combined to a single time trace. By applying the iterative *τ*_*c,opt*_ counting algorithm to the combined time traces as presented by Lee *et al.* (17) we finally derived the corresponding number of molecules per localization spot with minimal bias error.

#### Detectable fraction of PAmCherry molecules in E. coli

To extract a detectable fraction of PAmCherry for correction of LexA-PAmCherry counts, we expressed LacY-PAmCherry and LacY-tdPAmCherry from the original chromosomal locus in two *E. coli* strains. Well-separated signals were achieved by extremely low expression from *P*_*lac*_ using 17 µM IPTG. After single-molecule localization and counting as described above, we extracted the number of molecules per individual spot for each LacY construct. To reduce the error from non-matured or misfolded proteins, we restricted our analysis to a 200 nm wide region along the cell outline, which is where LacY localization is expected and predominately occurred when induced with a higher IPTG concentration of 150 µM (Fig. S5). As previously described (18), we built two histograms from the number of molecules per spot and applied a global binomial weighted least-square fit neglecting the first bin at *N* = 0 (Fig. S6). This resulted in a detectable fraction *F* = 0.479 ± 0.066 (one *σ* confidence interval).

#### Cell alignment

To create images of several cells stacked on top of each other to infer average localizations of LexA, HupA and LacY, we used the python package ColiCoords (71). First, *E. coli* outlines from non-dividing cells obtained using oufti were converted into binary images. These binaries, the corresponding brightfield images and PALM localization tables were then processed with ColiCoords functions to project localizations for all cells of a single condition into a synthetic cell with average dimensions. The resulting localization table was exported and visualized using the dscatter algorithm (72).

### Time course modeling

#### LexA synthesis, degradation model

The single-molecule counting time courses were modelled with a single population *V* (“visible”) for each stress condition. Synthesis of new molecules as well as the degradation of visible molecules over time were considered by *v*_*s*_ and 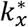 (Eq. 1, Fig. 5a).

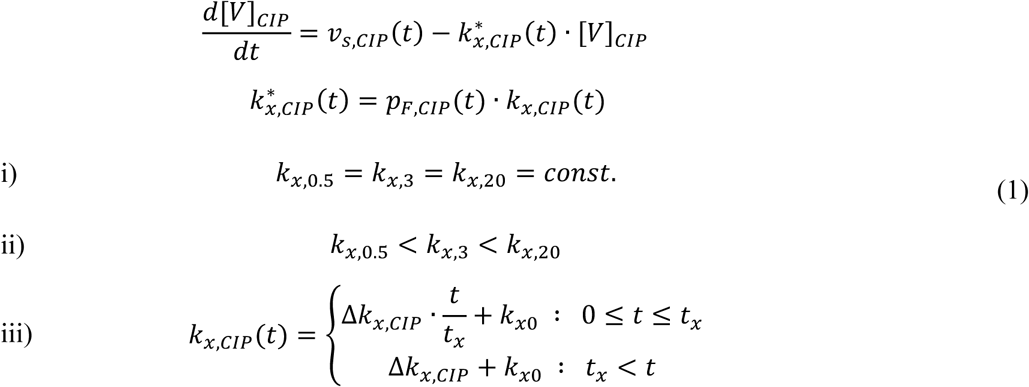

The time evolution of the concentration of visible molecules *V* was then fitted using numerical simulation with infinitesimal small time steps Δ*t*. For this purpose, Δ*t* was chosen such that the relative change of the concentration for each CIP condition was smaller than one, leading to Δ*t* = 0.01 min. The concentration of each time point was calculated iteratively by

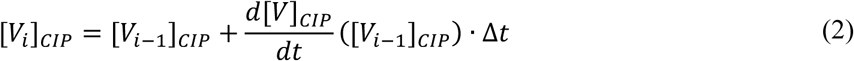

with [*V*_*i*_]_*CIP*_ being the concentration of visible molecules at time step *i* at the respective condition *CIP*. The time derivative, 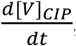, including synthesis and degradation is given in Eq. 1. The progressions of the measured concentrations at all stress conditions show a significant decrease followed by an increase of visible molecules, suggesting a time dependent *v*_*s*_(*t*). Since the time and the strength of the synthesis of new molecules appears to depend on the CIP concentration, and a maximal synthesis rate is expected, a logistic function was used to model *v*_*s*_(*t*):

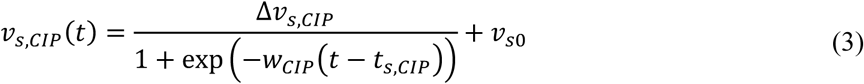

*v*_*s*0_ and Δ*v*_*s,CIP*_ are the synthesis speed at unstressed conditions and the maximum change of synthesis speed at the respective stress condition *CIP*. The corresponding response time and strength are described by *t*_*s,CIP*_ and *w*_*CIP*_. To stabilize the numerical fit, the following constraints have been used for the parameters of the response function.

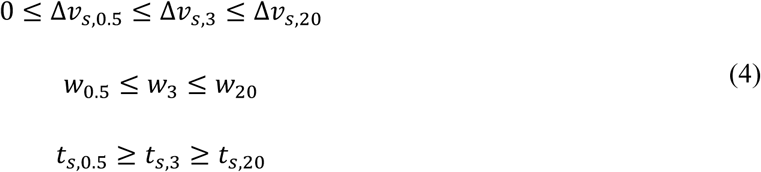

The apparent degradation rate 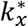, on the other hand, is defined as the product of the proportion of cleaved fragments *p*_*F,CIP*_(*t*) and the degradation rate *k*_*x,CIP*_(*t*) (Eq. 1). In order to disentangle the degradation rate from the product, the fractions of cleaved fragments derived by the single-molecule tracking measurements were fitted with a single exponential (Eq. 5) and incorporated in the counting fit.

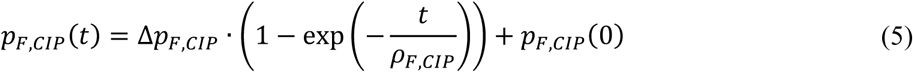

Here, *ρ*_*F,CIP*_ denotes the exponential time constant, and *p*_*F,CIP*_(0) and Δ*p*_*F,CIP*_ are the probability of cleaved fragments in absence of stress at *t* = 0 and the change of the probability over time, respectively.

Since the temporal behavior of *k*_*x,CIP*_ was unclear, three different scenarios were considered (Eq. 1):

i. A stress independent degradation rate, where all conditions and time points share the same rate.
ii. A time invariant but stress dependent degradation rate.
iii. A step function with shared zero stress rate and response time but increasing maximal degradation rate.

Fig. S6a shows the resulting *k*_*x,CIP*_(*t*) and *v*_*s*_(*t*) curves of global fits for the three scenarios. The corresponding fitting curves for the time evolution of visible molecules is exemplary shown for the stress condition [CIP] = 0.5 ng/ml for every scenario in the bottom panel. The fit quality was quantified by the reduced Chi-square 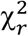, which also takes the number of fit parameters into account. The overall 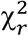 appears similar for the different scenarios with a value of around 20, however, the first case shows a significant overshoot in the beginning of the curve, followed by an increasing residual at larger *t*. The introduction of three different degradation rates in scenario ii) led to a decreased sum of squares of residuals for the cost of two more fitting parameters, indicating that *k*_*x*_ also depends on the concentration of CIP. Since an equal degradation rate is expected at zero stress, a shared degradation rate *k*_*x*0_ at *t* = 0 and saturation time *t*_*x*_ were introduced in scenario iii) leading to a smaller sum of squared residuals together with the reduction of the overshoot in the beginning of the curve.

Response time delays (Fig. 5g) were approximated from the difference of the characteristic time of an exponential fit of the first part of [*V*]_*CIP*_(*t*) and the actual response time of the respective *v*_*s,CIP*_(*t*) function.

#### Kinetic model for individual LexA populations

The LexA tracking data were modeled by four populations: target bound (T), DNA bound (D), cytoplasmic (C) and cleavage fragments (F). While LexA synthesis and degradation rates *v*_*s,CIP*_(*t*) and *k*_*x,CIP*_(*t*) were derived from the counting data, the interconversion between the individual populations is described by the rates *k*_*dc*_, *k*_*cd*_, *k*_*dt*_, *k*_*td*_, *k*_*tc*_, *k*_*ct*_, *k*_1_ (see Fig. 5b). As a first step, all rates were assumed to be independent on stress condition and constant over time. Global fitting of the time evolution of the four populations for all stress conditions was then performed by numerical simulation analogous to the counting fit (Eq. 2) with infinitesimal time steps of Δ*t* = 0.01 min. The corresponding time derivatives for each population are given in Eq. 6.

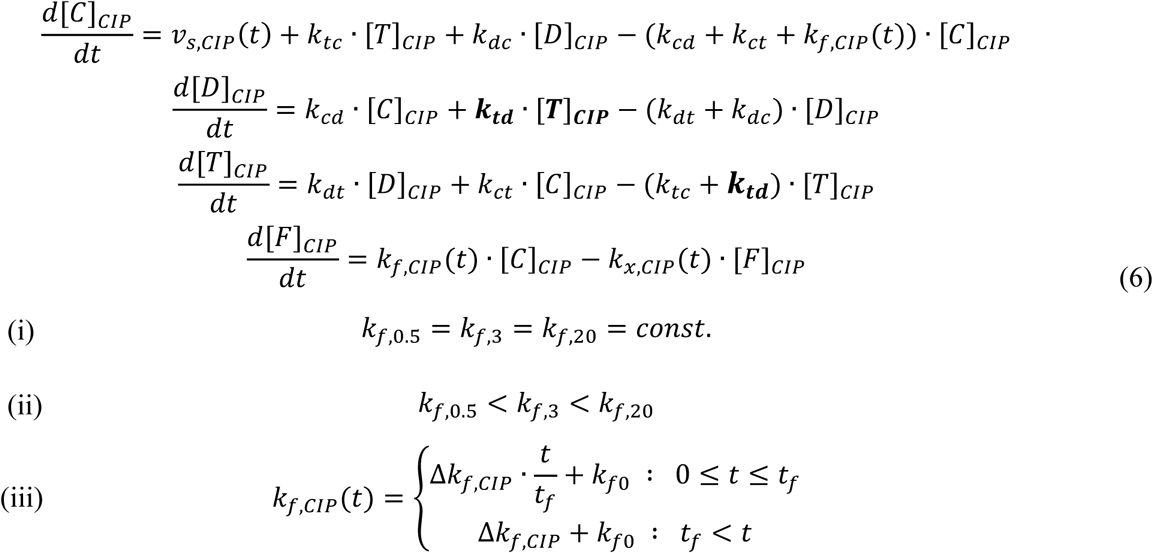

The fit result, exemplarily shown in Fig. S6b, led to a large residual with 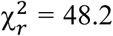. Since it is expected that the autoproteolysis of LexA depends on the inflicted DNA damage, individual cleavage rates *k*_*f,CIP*_ were used for the different CIP concentrations as a second scenario (Eq. 6ii). The corresponding fit led to a reduction of 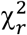 (Fig. S6b, mid panels). A shared cleavage rate at zero stress *k*_*f*0_ together with saturation time *t*_*f*_, were introduced, similar to the degradation rate. In this third scenario, 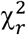 was further reduced. The final fit of scenario iii) also revealed that the interconversion rate from the target to the DNA bound state, *k*_*td*_, was close to zero and could therefore be neglected (bold in Eq. 6). In order to account for the random and systematic error of the extracted rates, bootstrapping was implemented in the fitting procedure, where for each time point of the tracking data a set of experimentally obtained *D*^∗^ histograms was drawn from the pool of biological replicates. The resulting rates and ratio of rates, *K*_*C*−*t*_ and *K*_*c*−*d*_, for each bootstrapping iteration are shown in Fig. 5e (gray) together with the best estimator of the combined replicates (black).

## Acknowledgements

We thank Frank Groß and Francis Stewart (TU Dresden) for recombineering plasmids and Thorsten Mascher (TU Dresden) for *E. coli* strain MG1655. We thank Edward Courchaine and Chris King (Yale University) for helpful comments on the manuscript. This research was supported by TU Dresden. We acknowledge support by Jens Ehrig of the Molecular Imaging and Manipulation Facility, a core facility of the CMCB at TU Dresden.

## Author Contributions

L.S. and M.S. designed the research, L.S. and M.T. carried out experiments, L.S. and A.H. analyzed data, A.H. implemented and developed algorithms and models, L.S. wrote the initial draft, L.S., M.T., A.H. and M.S. edited and contributed to the final manuscript, M.S. supervised the research and acquired funding.

## Supplementary Information

### LexA mainly localizes outside the nucleoid

Visual inspection of fixed-cell LexA PALM images suggested a seemingly uniform distribution of LexA in single cells (Fig. 6a). To reveal a possible preferential localization that might not be detectable in single cells due to too low LexA counts, we averaged LexA distributions over many cells (Fig. S5). We used separate strains with reference markers to determine LexA densities relative to the nucleoid (HU-PAmCherry) and membrane (LacY-PAmCherry). The three strains were imaged using PALM after fixation at 0, 60, 90 and 180 min with 3 ng/ml CIP stress. We used ColiCoords (71) to align localizations based on *E. coli* outlines from brightfield images (66). The average positioning of LacY along the cell periphery suggests a relatively high precision of this diffraction-limited method (Fig. S5). HU portrayed distributions within the cell center of unstressed cells in the region where the nucleoid is expected, in agreement with earlier reports (73, 74). Even under long stress, we observed a single, but diffuse DNA mass in the cell center. This is well in agreement with earlier reports for fluoroquinolone antibiotics (39, 75), but different from UV stress that can cause multiple segregated nucleoid-like structures in filamentous *E. coli* during late SOS response (40, 41). Interestingly, the average LexA^wt^ position does not colocalize with HU. In our 2D projection, LexA^wt^ explored a larger area, presumably including the cytoplasmic region between nucleoid and membrane. In fact, areas of particularly high LexA abundance appear outside of the high-density nucleoid regions. This was emphasized further throughout the SOS response and especially pronounced in images after 90 and 180 min CIP-stress. Such average localization agrees well with our single-molecule tracking analysis showing a high abundance of cytoplasmic and cleaved *vs.* DNA-associated repressors.

**Figure S1:**
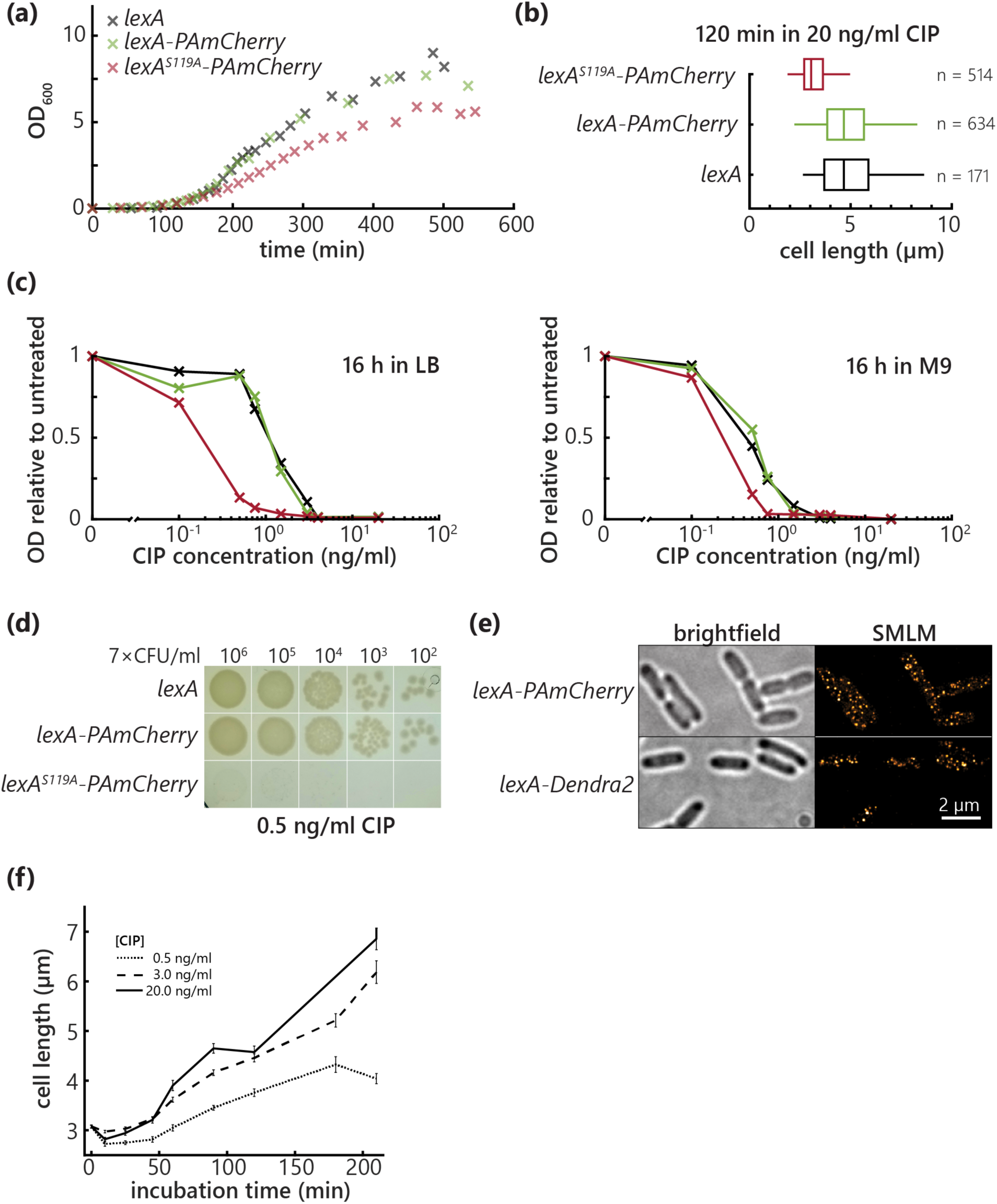
Growth assays and test of fusion protein function. **(a)** Growth curves of the three strains measured in M9 liquid medium. **(b)** Cell elongation during ciprofloxacin stress determined from outlines. Colors as in (a). **(c)** Growth capacity in LB medium during ciprofloxacin treatment in LB and M9 medium. Colors as in (a). **(d)** Agar plate growth assay on LB containing a low dose of ciprofloxacin. **(e)** Brightfield and single-molecule localization microscopy images of PFA-fixed *E. coli* strains having chromosomal *lexA* tagged with either PAmCherry or Dendra2. **(f)** Cell length increase during SOS response. Mean cell lengths at each time point are extracted from log-normal fits to length histograms. Error bars represent standard deviations from 50 bootstrapping iterations.

**Figure S2:**
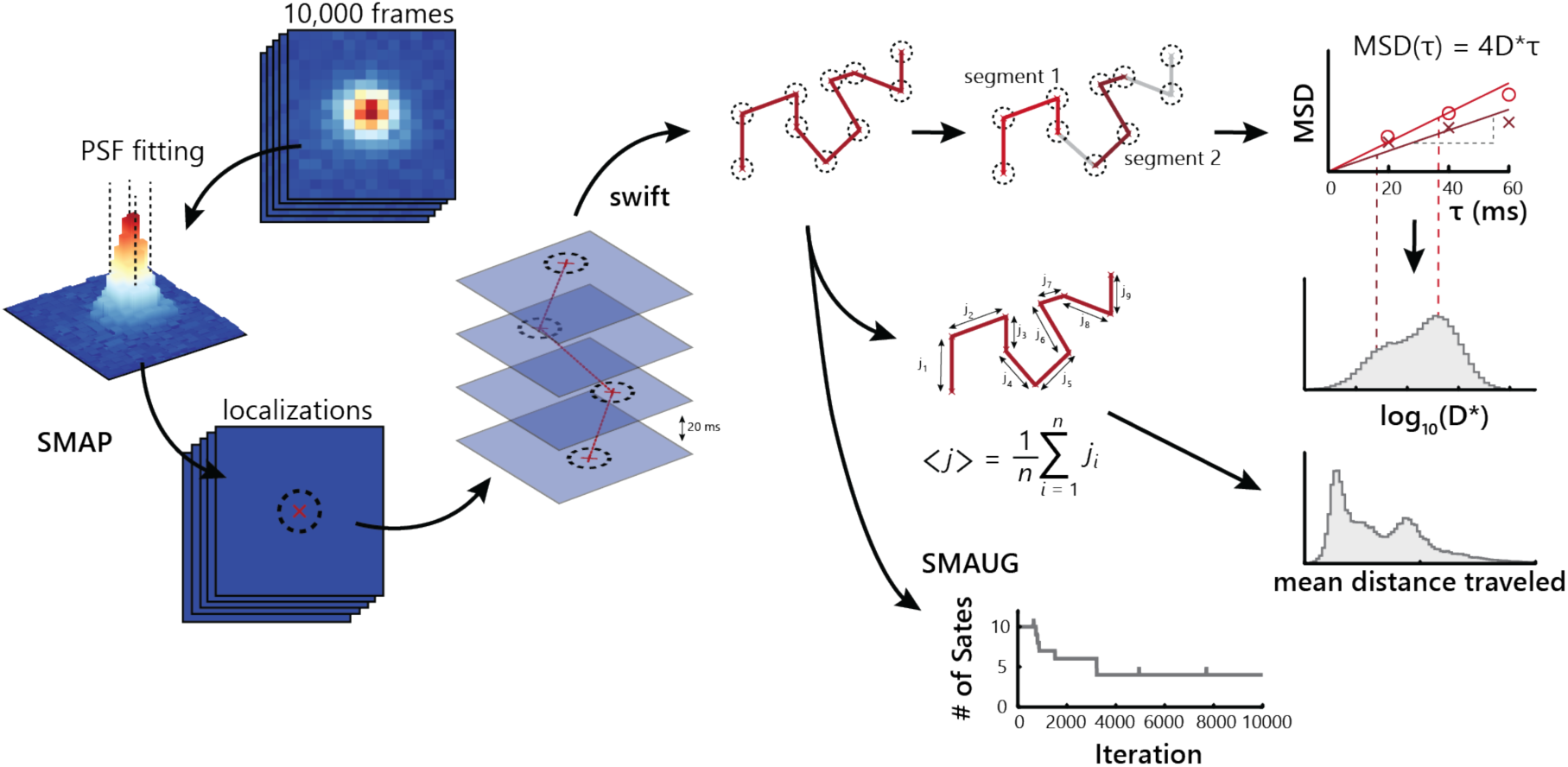
Single-molecule tracking analysis procedure. Raw PALM movies are processed with SMAP (22) to fit a PSF model to PAmCherry signals. The resulting localizations are then linked in time using the tracking software swift (23). Each trajectory can then be 1) analyzed with SMAUG or 2) subjected to a simple mean inter-frame-distance analysis or 3) split into four localizations long segments to determine apparent diffusion coefficients from linear mean squared displacement fits.

**Figure S3:**
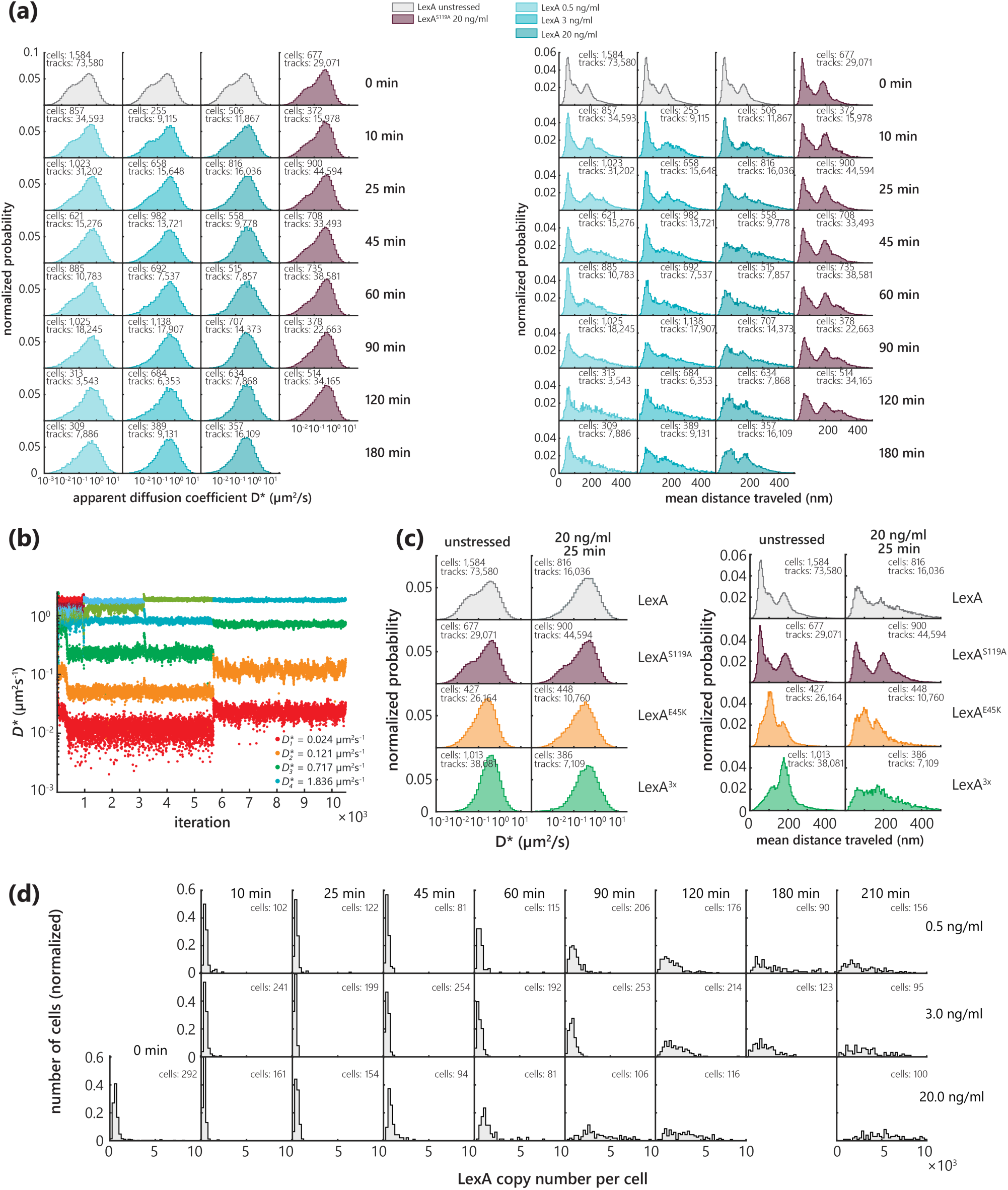
Single-molecule tracking and counting results for all experimental conditions. **(a)** Left: Apparent diffusion coefficient (*D\**) distributions of LexA-PAmCherry for all tracking experiments. Right: Histograms of mean distance traveled per frame of LexA-PAmCherry for all tracking experiments. Biological replicates are merged, cell and trajectory counts are indicated. **(b)** SMAUG analysis of LexA diffusion in stressed cells (20 ng/ml CIP, 45 min). **(c)** Left: Apparent diffusion coefficient (*D\**) distributions of LexA-PAmCherry mutants for all tracking experiments. Right: Histograms of mean distance traveled per frame of LexA-PAmCherry mutants for all tracking experiments. **(d)** LexA monomer copy number per cell. Counts are corrected for PAmCherry detectable fraction.

**Figure S4:**
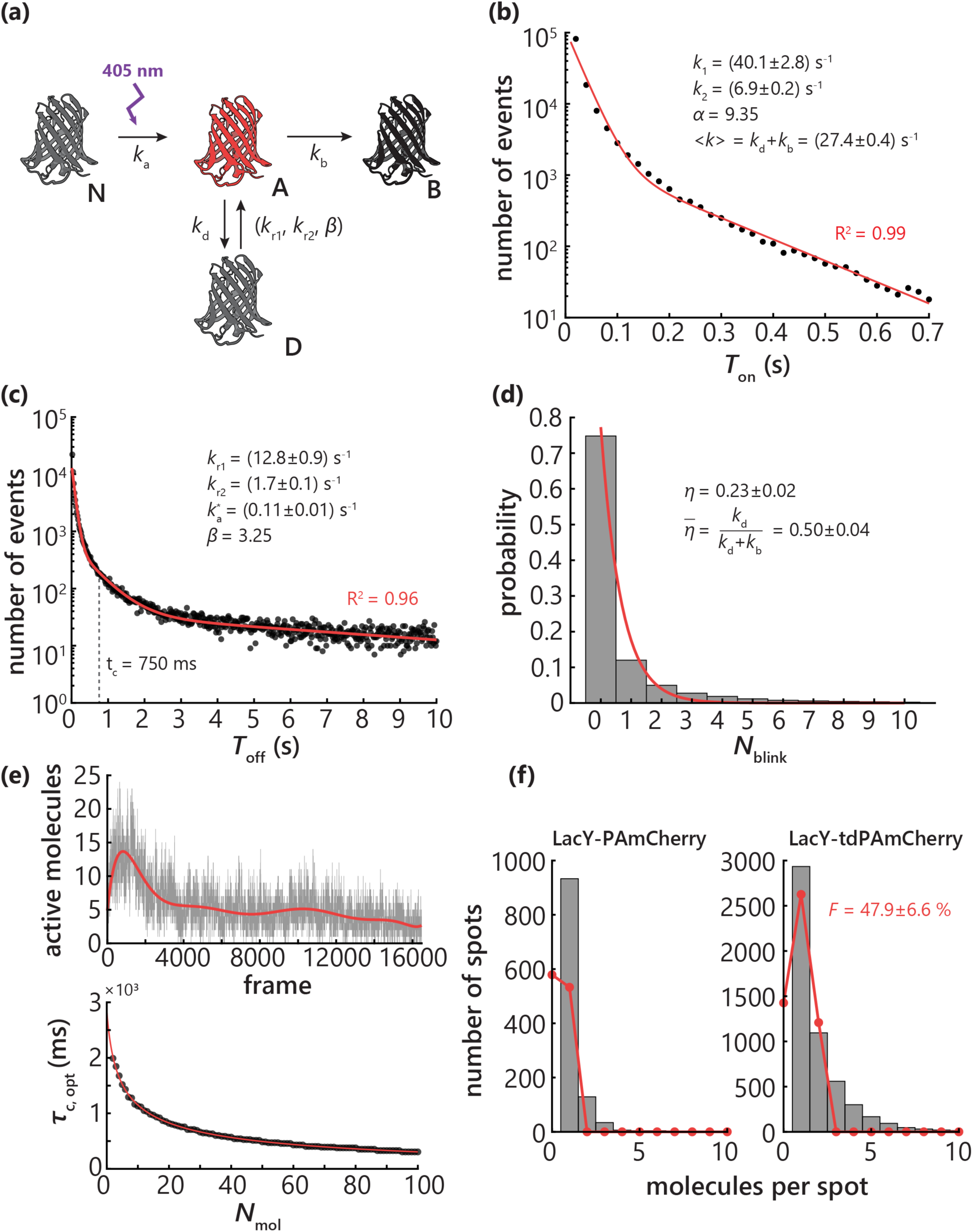
*In situ* Photophysical parameters for PAmCherry and single-molecule counting. **(a)** Reaction scheme for photophysical model of PAmCherry. **(b)** On-time distribution of PAmCherry in fixed *E. coli*. **(c)** Off-time distribution of PAmCherry in fixed *E. coli*. **(d)** Distribution of number of blinking events. **(e)** Top: Characteristic activation profile for a PALM movie (0.5 ng/ml CIP, 45 min), approximated with a polynomial fit. Bottom: Simulated optimal blinking tolerance time depending on the number of molecules in the time trace. **(f)** PAmCherry detectable fraction (*F*) determined from global binomial fits to distributions of number of molecules detected at the same spot for LacY-PAmCherry (left) and LacY-PAmCherry-PAmCherry (right).

**Figure S5:**
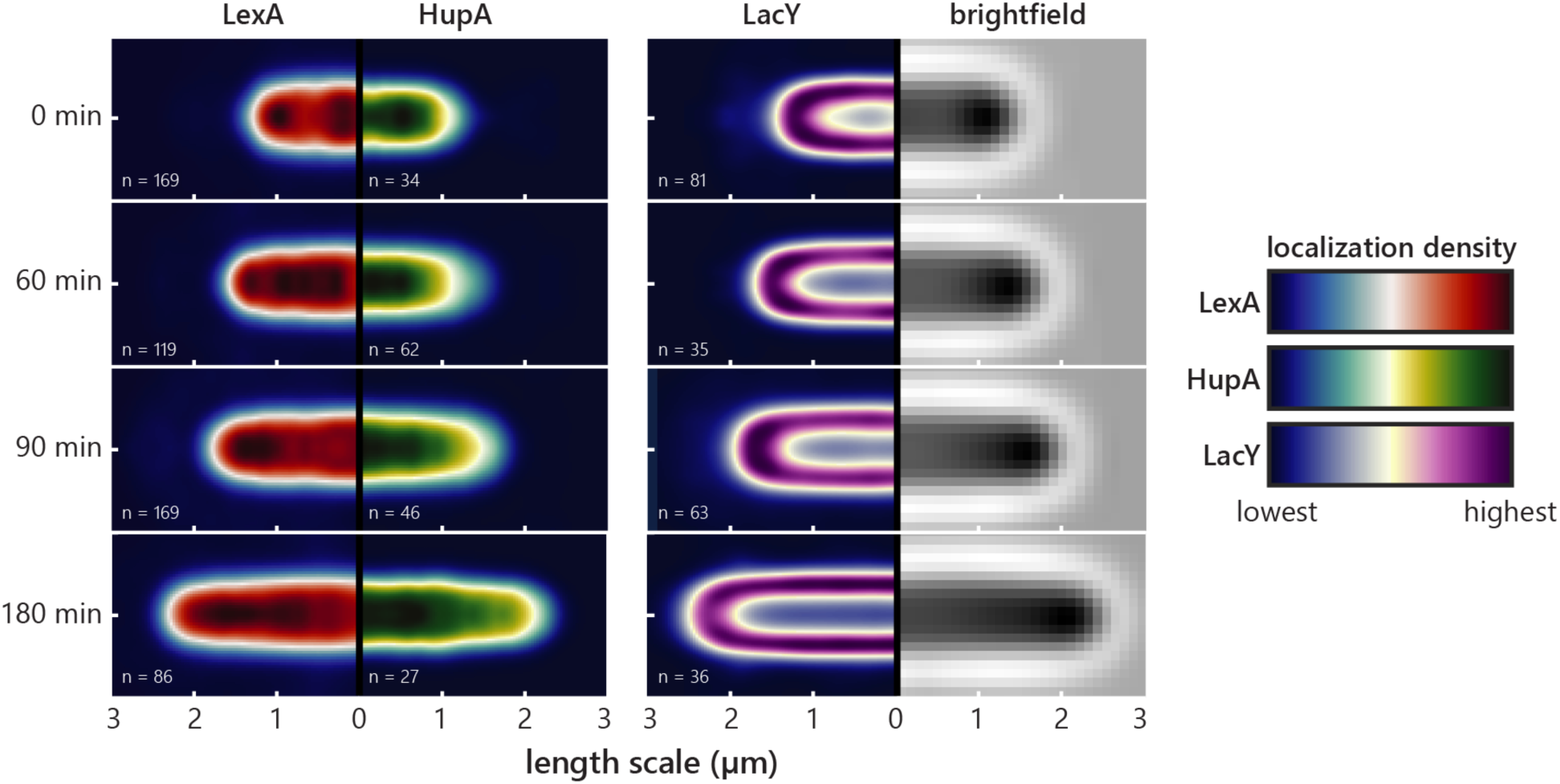
Average localization of LexA, HupA and LacY obtained by cell alignment. HupA-PAmCherry and LacY-PAmCherry serve as nucleoid and membrane-markers, respectively. Dividing cells and cells with lengths outside the standard deviation of the associated population were excluded. Localizations from *n* = 27-169 cells were projected into a synthetic cell with average dimensions. Incubation times with 1×MIC CIP are indicated on the left.

**Figure S6:**
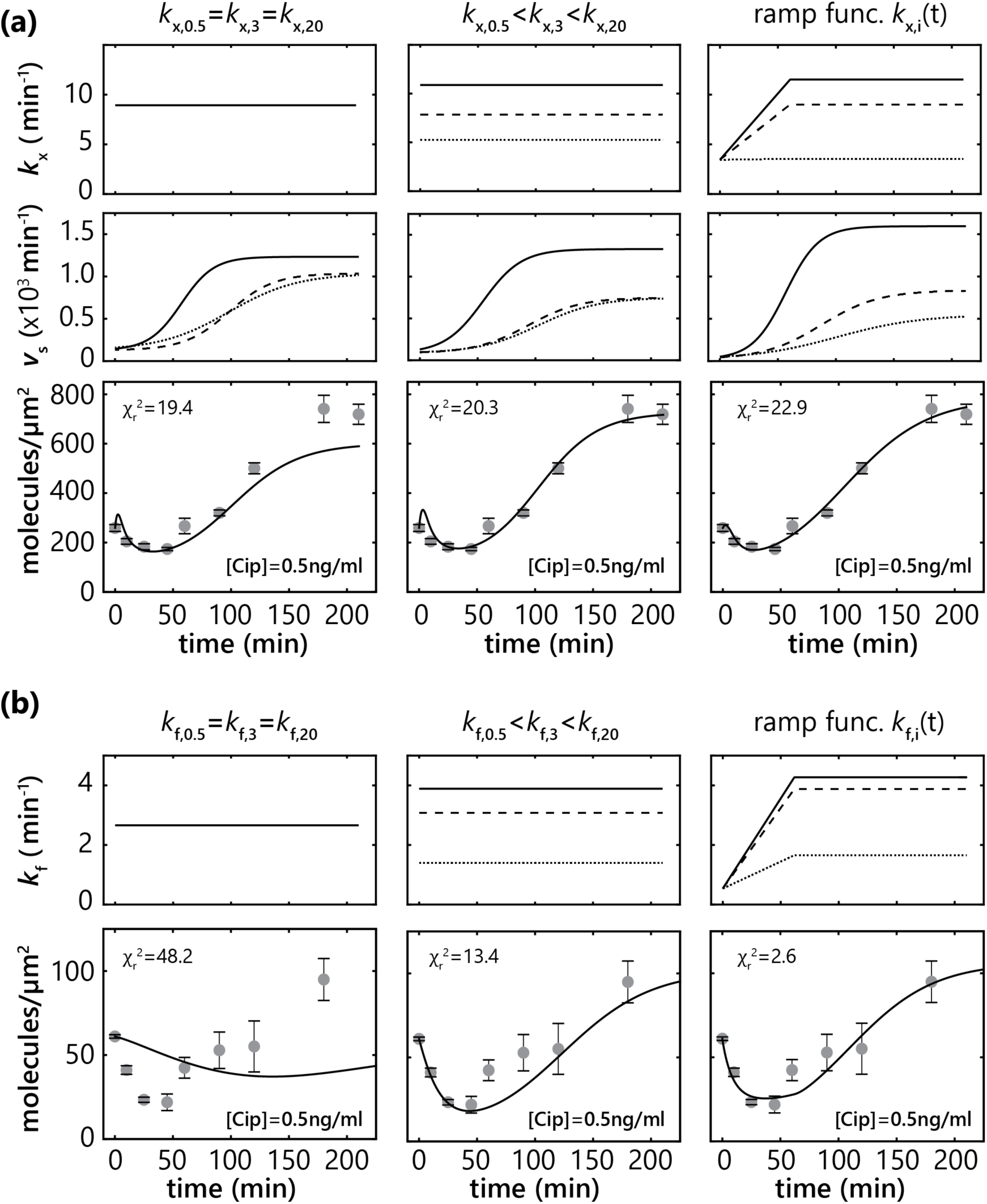
Kinetic model development. Different scenarios for the degradation rate k_x_ incorporated in the fitting of the single-molecule counting data **(a)** and for the fragmentation rate k_f_ incorporated in the fitting of the single-molecule tracking data **(b)**.

**Tab. S1:** Photophysical parameters for PAmCherry in *E. coli*.

| $k_1$ (s<sup>-1</sup>) | $k_2$ (s<sup>-1</sup>) | $\alpha$ | $\langle k \rangle$ (s<sup>-1</sup>) | $k_{r1}$ (s<sup>-1</sup>) | $k_{r2}$ (s<sup>-1</sup>) | $\beta$ |
| --- | --- | --- | --- | --- | --- | --- |
| 40.1±2.8 | 6.9±0.2 | 9.35 | 27.4±0.4 | 12.8±0.9 | 1.7±0.1 | 3.25 |
| $\eta$ | $\bar{\eta}$ | $k_b$ (s<sup>-1</sup>) | $k_d$ (s<sup>-1</sup>) | $p_{\text{reappear}}$ | $p_{\text{bleach}}$ | $p_{\text{blink}}$ |
| 0.23±0.02 | 0.50±0.04 | 13.70±1.06 | 13.71±1.05 | 0.1017 | 0.2739 | 0.2741 |

**Tab. S2:**
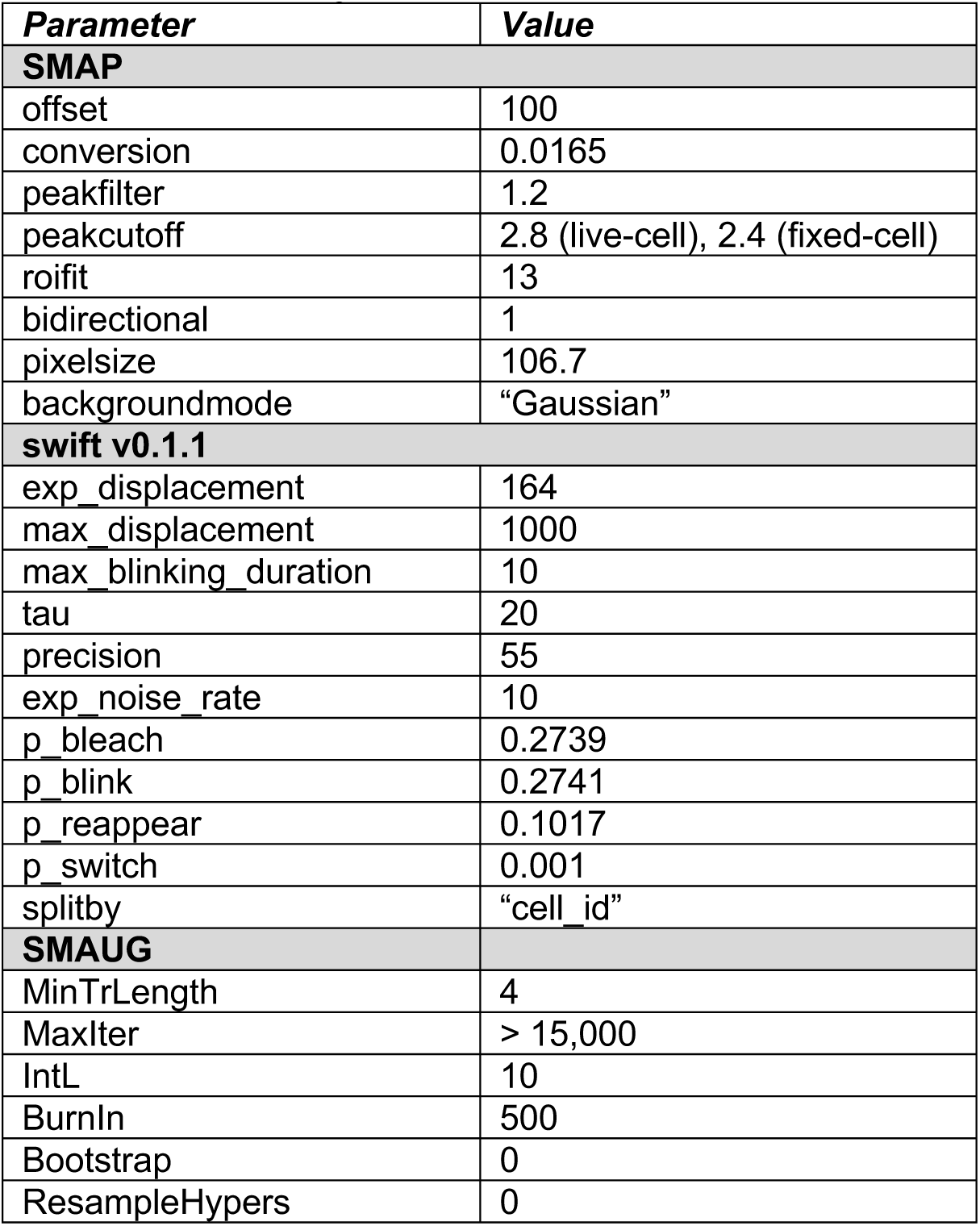
Parameters for algorithms.

| **Parameter** | **Value** |
| --- | --- |
| **SMAP** |  |
| offset | 100 |
| conversion | 0.0165 |
| peakfilter | 1.2 |
| peakcutoff | 2.8 (live-cell), 2.4 (fixed-cell) |
| roifit | 13 |
| bidirectional | 1 |
| pixelsize | 106.7 |
| backgroundmode | "Gaussian" |
| **swift v0.1.1** |  |
| exp_displacement | 164 |
| max_displacement | 1000 |
| max_blinking_duration | 10 |
| tau | 20 |
| precision | 55 |
| exp_noise_rate | 10 |
| p_bleach | 0.2739 |
| p_blink | 0.2741 |
| p_reappear | 0.1017 |
| p_switch | 0.001 |
| splitby | "cell_id" |
| **SMAUG** |  |
| MinTrLength | 4 |
| MaxIter | > 15,000 |
| IntL | 10 |
| BurnIn | 500 |
| Bootstrap | 0 |
| ResampleHypers | 0 |

## References

1. H. Nikaido, Multidrug Resistance in Bacteria. Annu. Rev. Biochem. 78, 119–146 (2009).

2. Z. Baharoglu, D. Mazel, SOS, the formidable strategy of bacteria against aggressions. FEMS Microbiol. Rev. 38, 1126–1145 (2014).

3. L. A. Simmons, J. J. Foti, S. E. Cohen, G. C. Walker, The SOS Regulatory Network. EcoSal Plus, 1–30 (2008).

4. M. Butala, D. Žgur-Bertok, S. J. W. Busby, The bacterial LexA transcriptional repressor. Cell. Mol. Life Sci. 66, 82–93 (2009).

5. C. Joo, et al., Real-Time Observation of RecA Filament Dynamics with Single Monomer Resolution. Cell 126, 515–527 (2006).

6. M. Sassanfar, J. W. Roberts, Nature of the SOS-inducing signal in Escherichia coli. J. Mol. Biol. 212, 79–96 (1990).

7. J. W. Little, LexA cleavage and other self-processing reactions. J. Bacteriol. 175, 4943–50 (1993).

8. M. Butala, et al., Interconversion between bound and free conformations of LexA orchestrates the bacterial SOS response. Nucleic Acids Res. 39, 6546–6557 (2011).

9. M. J. Culyba, J. M. Kubiak, C. Y. Mo, M. Goulian, R. M. Kohli, Non-equilibrium repressor binding kinetics link DNA damage dose to transcriptional timing within the SOS gene network. PLoS Genet. 14, 1–29 (2018).

10. S. B. Neher, J. M. Flynn, R. T. Sauer, T. A. Baker, Latent ClpX-recognition signals ensure LexA destruction after DNA damage. Genes Dev. 17, 1084–9 (2003).

11. N. Friedman, S. Vardi, M. Ronen, U. Alon, J. Stavans, Precise temporal modulation in the response of the SOS DNA repair network in individual bacteria. PLoS Biol. 3, 1261–1268 (2005).

12. Y. Shimoni, S. Altuvia, H. Margalit, O. Biham, Stochastic analysis of the SOS response in Escherichia coli. PLoS One 4 (2009).

13. J. D. McCool, et al., Measurement of SOS expression in individual Escherichia coli K-12 cells using fluorescence microscopy. Mol. Microbiol. 53, 1343–1357 (2004).

14. J. Elf, I. Barkefors, Single-Molecule Kinetics in Living Cells. Annu. Rev. Biochem. 88, 635–659 (2019).

15. A. N. Kapanidis, A. Lepore, M. El Karoui, Rediscovering Bacteria through Single-Molecule Imaging in Living Cells. Biophys. J. (2018) https://doi.org/10.1016/j.bpj.2018.03.028.

16. I. Vojnovic, J. Winkelmeier, U. Endesfelder, Visualizing the inner life of microbes: practices of multi-color single-molecule localization microscopy in microbiology. Biochem. Soc. Trans. 47, 1041–1065 (2019).

17. S.-H. Lee, J. Y. Shin, A. Lee, C. Bustamante, Counting single photoactivatable fluorescent molecules by photoactivated localization microscopy (PALM). Proc. Natl. Acad. Sci. 109, 17436–17441 (2012).

18. E. M. Puchner, J. M. Walter, R. Kasper, B. Huang, W. A. Lim, Counting molecules in single organelles with superresolution microscopy allows tracking of the endosome maturation trajectory. Proc. Natl. Acad. Sci. 110, 16015–16020 (2013).

19. S. Wang, J. R. Moffitt, G. T. Dempsey, X. S. Xie, X. Zhuang, Characterization and development of photoactivatable fluorescent proteins for single-molecule-based superresolution imaging. Proc. Natl. Acad. Sci. 111, 8452–8457 (2014).

20. A. P. P. Zhang, Y. Z. Pigli, P. A. Rice, Structure of the LexA-DNA complex and implications for SOS box measurement. Nature 466, 883–886 (2010).

21. D. Landgraf, B. Okumus, P. Chien, T. A. Baker, J. Paulsson, Segregation of molecules at cell division reveals native protein localization. Nat. Methods 9, 480–482 (2012).

22. Y. Li, et al., Real-time 3D single-molecule localization using experimental point spread functions. Nat. Methods 15, 367–369 (2018).

23. M. Endesfelder, C. Schießl, B. Turkowyd, T. Lechner, U. Endesfelder, swift – fast, probabilistic tracking for dense, highly dynamic single-molecule data. Manuscr. Prep. (2020).

24. B. M. A. Howard, R. J. Pinney, J. T. Smith, Function of the SOS Process in Repair of DNA Damage Induced by Modern 4-Quinolones. J. Pharm. Pharmacol. 45, 658–662 (1993).

25. P. Wang, et al., Subinhibitory concentrations of ciprofloxacin induce SOS response and mutations of antibiotic resistance in bacteria. Ann. Microbiol. 60, 511–517 (2010).

26. K. Drlica, M. Malik, R. J. Kerns, X. Zhao, Quinolone-Mediated Bacterial Death. Antimicrob. Agents Chemother. 52, 385–392 (2008).

27. J. D. Karslake, et al., SMAUG: Analyzing single-molecule tracks with nonparametric Bayesian statistics. Methods (2020) https://doi.org/10.1016/j.ymeth.2020.03.008.

28. A. T. Thliveris, D. W. Mount, Genetic identification of the DNA binding domain of Escherichia coli LexA protein. Proc. Natl. Acad. Sci. 89, 4500–4504 (1992).

29. S. N. Slilaty, J. W. Little, Lysine-156 and serine-119 are required for LexA repressor cleavage: a possible mechanism. Proc. Natl. Acad. Sci. 84, 3987–3991 (1987).

30. L. Schärfen, M. Tišma, M. Schlierf, Fast, simultaneous tagging and mutagenesis of genes on bacterial chromosomes. ACS Synth. Biol. (2020) https://doi.org/10.1021/acssynbio.0c00202.

31. L. J. Goodman, R. M. Fliegelman, G. M. Trenholme, R. L. Kaplan, Comparative in vitro activity of ciprofloxacin against Campylobacter spp. and other bacterial enteric pathogens. Antimicrob. Agents Chemother. 25, 504–506 (1984).

32. J. M. Andrews, Determination of minimum inhibitory concentrations. J. Antimicrob. Chemother. 48, 5–16 (2001).

33. G. V. Smirnova, O. N. Oktyabrsky, Relationship between Escherichia coli growth rate and bacterial susceptibility to ciprofloxacin. FEMS Microbiol. Lett. 365, 1–6 (2018).

34. A. Gutierrez, et al., Understanding and Sensitizing Density-Dependent Persistence to Quinolone Antibiotics. Mol. Cell 68, 1147-1154.e3 (2017).

35. H. Deschout, A. Shivanandan, P. Annibale, M. Scarselli, A. Radenovic, Progress in quantitative single-molecule localization microscopy. Histochem. Cell Biol. 142, 5–17 (2014).

36. N. Durisic, L. Laparra-Cuervo, Á. Sandoval-Álvarez, J. S. Borbely, M. Lakadamyali, Single-molecule evaluation of fluorescent protein photoactivation efficiency using an in vivo nanotemplate. Nat. Methods 11, 156–162 (2014).

37. F. Fricke, J. Beaudouin, R. Eils, M. Heilemann, One, two or three? Probing the stoichiometry of membrane proteins by single-molecule localization microscopy. Sci. Rep. 5, 1–8 (2015).

38. A. J. B. Kreutzberger, S. Urban, Single-Molecule Analyses Reveal Rhomboid Proteins Are Strict and Functional Monomers in the Membrane. Biophys. J. 115, 1755–1761 (2018).

39. J. Cambridge, A. Blinkova, D. Magnan, D. Bates, J. R. Walker, A Replication-Inhibited Unsegregated Nucleoid at Mid-Cell Blocks Z-Ring Formation and Cell Division Independently of SOS and the SlmA Nucleoid Occlusion Protein in Escherichia coli. J. Bacteriol. 196, 36–49 (2014).

40. I. Odsbu, K. Skarstad, DNA compaction in the early part of the SOS response is dependent on RecN and RecA. Microbiology 160, 872–882 (2014).

41. J. Cayron, A. Dedieu, C. Lesterlin, Bacterial filament division dynamics allows rapid post-stress cell proliferation. BioRxiv (2020).

42. J. Hettich, J. C. M. Gebhardt, Transcription factor target site search and gene regulation in a background of unspecific binding sites. J. Theor. Biol. 454, 91–101 (2018).

43. A. Sanchez, I. Golding, Genetic determinants and cellular constraints in noisy gene expression. Science 342, 1188–93 (2013).

44. F. L. M. Vallania, et al., Origin and Consequences of the Relationship between Protein Mean and Variance. PLoS One 9, e102202 (2014).

45. Y. Taniguchi, et al., Quantifying E. coli proteome and transcriptome with single-molecule sensitivity in single cells. Science 329, 533–8 (2010).

46. J. Courcelle, A. Khodursky, B. Peter, P. O. Brown, P. C. Hanawalt, Comparative gene expression profiles following UV exposure in wild-type and SOS-deficient Escherichia coli. Genetics 158, 41–64 (2001).

47. P. Hammar, et al., The lac repressor displays facilitated diffusion in living cells. Science 336, 1595–8 (2012).

48. F. Garza de Leon, L. Sellars, M. Stracy, S. J. W. Busby, A. N. Kapanidis, Tracking Low-Copy Transcription Factors in Living Bacteria: The Case of the lac Repressor. Biophys. J. 112, 1316–1327 (2017).

49. A. Javer, et al., Short-time movement of E. coli chromosomal loci depends on coordinate and subcellular localization. Nat. Commun. 4, 3003 (2013).

50. Y. Zhu, S. Mohapatra, J. C. Weisshaar, Rigidification of the Escherichia coli cytoplasm by the human antimicrobial peptide LL-37 revealed by superresolution fluorescence microscopy. Proc. Natl. Acad. Sci. 116, 1017–1026 (2019).

51. S. Hegde, S. J. Sandler, A. J. Clark, M. V. V. S. Madiraju, recO and recR mutations delay induction of the SOS response in Escherichia coli. MGG Mol. Gen. Genet. 246, 254–258 (1995).

52. S. V Aksenov, Induction of the SOS Response in Ultraviolet-Irradiated Escherichia coli Analyzed by Dynamics of LexA, RecA and SulA Proteins. J. Biol. Phys. 25, 263–277 (1999).

53. A. N. Bugay, E. A. Krasavin, A. Y. Parkhomenko, M. A. Vasilyeva, Modeling nucleotide excision repair and its impact on UV-induced mutagenesis during SOS-response in bacterial cells. J. Theor. Biol. 364, 7–20 (2015).

54. K. C. Giese, C. B. Michalowski, J. W. Little, RecA-Dependent Cleavage of LexA Dimers. J. Mol. Biol. 377, 148–161 (2008).

55. K. L. Roland, M. H. Smith, J. A. Rupley, J. W. Little, In vitro analysis of mutant LexA proteins with an increased rate of specific cleavage. J. Mol. Biol. 228, 395–408 (1992).

56. J. Winkler, et al., Quantitative and spatio-temporal features of protein aggregation in Escherichia coli and consequences on protein quality control and cellular ageing. EMBO J. 29, 910–923 (2010).

57. S. Bakshi, A. Siryaporn, M. Goulian, J. C. Weisshaar, Superresolution imaging of ribosomes and RNA polymerase in live Escherichia coli cells. Mol. Microbiol. 85, 21–38 (2012).

58. J. Yu, Probing Gene Expression in Live Cells, One Protein Molecule at a Time. Science 311, 1600–1603 (2006).

59. N. Dhar, J. D. McKinney, Microbial phenotypic heterogeneity and antibiotic tolerance. Curr. Opin. Microbiol. 10, 30–38 (2007).

60. S. Uphoff, Real-time dynamics of mutagenesis reveal the chronology of DNA repair and damage tolerance responses in single cells. Proc. Natl. Acad. Sci. 115, E6516–E6525 (2018).

61. F. Goormaghtigh, L. Van Melderen, Single-cell imaging and characterization of Escherichia coli persister cells to ofloxacin in exponential cultures. Sci. Adv. 5, eaav9462 (2019).

62. T. C. Barrett, W. W. K. Mok, A. M. Murawski, M. P. Brynildsen, Enhanced antibiotic resistance development from fluoroquinolone persisters after a single exposure to antibiotic. Nat. Commun. 10, 1177 (2019).

63. Y. Zhang, F. Buchholz, J. P. P. Muyrers, A. F. Stewart, A new logic for DNA engineering using recombination in Escherichia coli. Nat. Genet. 20, 123–128 (1998).

64. H. Wang, et al., RecET direct cloning and Redαβ recombineering of biosynthetic gene clusters, large operons or single genes for heterologous expression. Nat. Protoc. 11, 1175–1190 (2016).

65. S. O. Skinner, L. A. Sepúlveda, H. Xu, I. Golding, Measuring mRNA copy number in individual Escherichia coli cells using single-molecule fluorescent in situ hybridization. Nat. Protoc. 8, 1100–1113 (2013).

66. A. Paintdakhi, et al., Oufti: An integrated software package for high-accuracy, high-throughput quantitative microscopy analysis. Mol. Microbiol. 99, 767–777 (2016).

67. M. Vrljic, S. Y. Nishimura, S. Brasselet, W. E. Moerner, H. M. McConnell, Translational Diffusion of Individual Class II MHC Membrane Proteinsin Cells. Biophys. J. 83, 2681–2692 (2002).

68. A. Balinovic, D. Albrecht, U. Endesfelder, Spectrally red-shifted fluorescent fiducial markers for optimal drift correction in localization microscopy. J. Phys. D. Appl. Phys. 52, 204002 (2019).

69. T. Pengo, S. J. Holden, S. Manley, PALMsiever: A tool to turn raw data into results for single-molecule localization microscopy. Bioinformatics 31, 797–798 (2015).

70. M. Ovesný, P. Křížek, J. Borkovec, Z. Švindrych, G. M. Hagen, ThunderSTORM: A comprehensive ImageJ plug-in for PALM and STORM data analysis and super-resolution imaging. Bioinformatics 30, 2389–2390 (2014).

71. J. H. Smit, Y. Li, E. M. Warszawik, A. Herrmann, T. Cordes, ColiCoords: A Python package for the analysis of bacterial fluorescence microscopy data. PLoS One 14, e0217524 (2019).

72. P. H. C. Eilers, J. J. Goeman, Enhancing scatterplots with smoothed densities. Bioinformatics 20, 623–628 (2004).

73. C. Spahn, F. Cella-Zannacchi, U. Endesfelder, M. Heilemann, Correlative super-resolution imaging of RNA polymerase distribution and dynamics, bacterial membrane and chromosomal structure in Escherichia coli. Methods Appl. Fluoresc. 3 (2015).

74. M. Stracy, et al., Single-molecule imaging of UvrA and UvrB recruitment to DNA lesions in living Escherichia coli. Nat. Commun. 7, 1–9 (2016).

75. N. H. Georgopapadakou, A. Bertasso, Effects of quinolones on nucleoid segregation in Escherichia coli. Antimicrob. Agents Chemother. 35, 2645–2648 (1991).

